# Physics-based inverse design of cholesterol attracting transmembrane helices reveals a paradoxical role of hydrophobic length

**DOI:** 10.1101/2021.07.01.450699

**Authors:** Jeroen Methorst, Nino Verwei, Christian Hoffmann, Paweł Chodnicki, Roberto Sansevrino, Han Wang, Niek van Hilten, Dennis Aschmann, Alexander Kros, Loren Andreas, Jacek Czub, Dragomir Milovanovic, Herre Jelger Risselada

**Affiliations:** Leiden Institute of Chemistry, Leiden University, The Netherlands; Department of Physics, Technical University of Dortmund, Germany; Department of Physical Chemistry, Gdansk university of technology, Poland; German Center for Neurodegenerative Diseases (DZNE), Research Site at Charité - Universitätsmedizin Berlin; Department of NMR-based Structural Biology, Max Planck Institute for Multidisciplinary Sciences, Göttingen, Germany

**Keywords:** genetic algorithm, molecular dynamics, artificial intelligence, Evo-MD, physics-based inverse design

## Abstract

The occurrence of linear cholesterol-recognition motifs in alpha-helical transmembrane domains has long been debated. Here, we demonstrate the ability of a genetic algorithm guided by coarse-grained molecular dynamics simulations—a method coined evolutionary molecular dynamics (Evo-MD)—to directly resolve the sequence which maximally attracts cholesterol for single-pass alpha-helical transmembrane domains (TMDs). We illustrate that the evolutionary landscape of cholesterol attraction in membrane proteins is characterized by a sharp, well-defined global optimum. Surprisingly, this optimal solution features an unusual short, slender hydrophobic block surrounded by three successive lysines. Owing to the membrane thickening effect of cholesterol, cholesterol-enriched ordered phases favor TMDs characterized by a long rather than a too short hydrophobic length (a negative hydrophobic mismatch). However, this short hydrophobic pattern evidently offers a pronounced net advantage for the attraction of free cholesterol in both coarse-grained and atomistic simulations. We illustrate that optimal cholesterol attraction is in fact based on the superposition of two distinct structural features: (i) slenderness and (ii) hydrophobic mismatch. In addition, we explore the evolutionary occurrence and feasibility of the two features by analyzing existing databases of membrane proteins and through the direct expression of analogous short hydrophobic sequences in live cell assays. The puzzling sequence variability of proposed linear cholesterol-recognition motifs is indicative of a sub-optimal membrane-mediated attraction of cholesterol which markedly differs from ligand binding based on shape compatibility.

**Significance Statement:** Our work demonstrates how a synergy between evolutionary algorithms and high-throughput coarse-grained molecular dynamics can yield fundamentally new insights into the evolutionary fingerprints of protein-mediated lipid sorting. We illustrate that the evolutionary landscape of cholesterol attraction in isolated transmembrane domains is characterized by a well-defined global optimum. In contrast, sub-optimal attraction of cholesterol is associated with a diverse solution space and features a high sequence variability despite acting on the same unique molecule. The contrasting physicochemical nature of the resolved attraction optimum suggests that cholesterol attraction via linear motifs does not pose a dominant pressure on the evolution of transmembrane proteins.

---

**C**holesterol serves as a major constituent of the mammalian plasma membrane. The overall fraction of cholesterol in the plasma membrane relative to total plasma membrane lipids is about 30% to 40% in leukocytes, epithelial cells, neurons, and mesenchymal cells (1). The localization, trafficking, and functionality of membrane proteins involved in cholesterol-dependent pathways and cholesterol homeostasis may critically rely on their ability to attract and bind cholesterol molecules (2–10). Prediction of protein-cholesterol affinity could therefore illuminate their role in diseases that are characterized by loss of cholesterol homeostasis (e.g. neurological diseases and cancer (11)), and pave the road for novel drug targets and strategies (6, 12–16). A compelling amount of data obtained by bioinformatic approaches, molecular modeling and simulations, and experiments have suggested the existence of cholesterol recognition amino acid consensus motifs (CRAC motifs) (3, 4, 17, 18), as well as its inverse (CARC motif), in various membrane protein families, including, for example: viral membrane proteins (e.g. (12, 15)), ion channels (e.g. (19, 20)), and G protein-coupled receptors (GPCRs)— the most intensively studied drug target family (e.g. (6, 21–24)). However, the looseness of the CRAC definition, being a rather flexible algorithmic rule: (L/V)-X_1*−*5_-(Y)-X_1*−*5_-(K/R), is rather unexpected for a motif that mediates binding to a unique molecule, raising skepticism about its predictive value (3). Interestingly, a recent fluorescence microscopy study on model membranes suggested that ‘slender’ transmembrane domains (TMDs) comprised of hydrophobic amino acids with short side chains, such as leucine (L) or valine (V), preferentially bind to the cholesterol-enriched interface of liquid order (Lo) domains (5). These amino acids also play a vital role in the CRAC motif. The observation that partitioning towards cholesterol-enriched phases strongly correlates to the accessible surface area of TMDs suggests that cholesterol recognition is not mediated by precise matching of molecular shape compatibility, as is common in protein-ligand docking, but is likely additionally mediated by alternative thermodynamic driving forces. This would explain why these amino acids occur on variable positions within the CRAC definition.

## Evolutionary Molecular Dynamics (Evo-MD)

High-throughput screening of transmembrane sequences could provide an effective strategy to resolve the chemical and structural specificities which underpin cholesterol recognition and binding, as well as the relevant thermodynamic driving forces. However, the accessible chemical space of transmembrane domains is astronomical (about 20^20^ possibilities), warranting the employment of smart search strategies.

Directed evolution is a method used in protein engineering that mimics the process of natural selection to steer proteins or nucleic acids toward a user-specified goal(25). Evolutionary inverse design strategies see applications in a variety of fields due to their efficient exploration of search-space (26). These methods fall under the scope of reinforcement learning, adapting processes for optimal performance by reinforcing desired behavior (27). Of special interest are the genetic algorithms (GA), which model the mechanisms of Darwinistic evolution in a computational algorithm, utilizing genetic elements such as recombination, cross-over, mutation, selection, and fitness (28). Since directed evolution is both time and labor intensive, it can quickly become intractable in a laboratory setting thereby limiting its application. In such scenarios, molecular dynamics (MD) simulations may provide an alternate *in silico* route for the high-throughput virtual screening of chemical space.

Here, we demonstrate the ability of GAs guided by coarse-grained MD simulations—a method which we coin evolutionary molecular dynamics (Evo-MD)—to yield unique insights into the driving forces that underpin ligand recognition (see Fig. 1). Evo-MD effectively reduces the search for optimal ligand consensus motifs to solving a variational problem in high dimensional chemical space. While many implementations of GAs have had abstract purposes—often used as a mathematical rather than a biological approach—we effectively create an *in silico* molecular cell membrane environment subject to natural evolution. To this aim, we introduce EVO-MD, a highly parallel software package for evolutionary molecular dynamics simulations that incorporates the GROMACS molecular dynamics engine into a custom, Python-based GA wrapper. EVO-MD can adapt any element of MD simulations, be it structural (e.g. atoms, molecules), topological, or simulation parameters (e.g. forcefield parameters), based on a reinforcement value measured during the simulation.

**Fig. 1.**
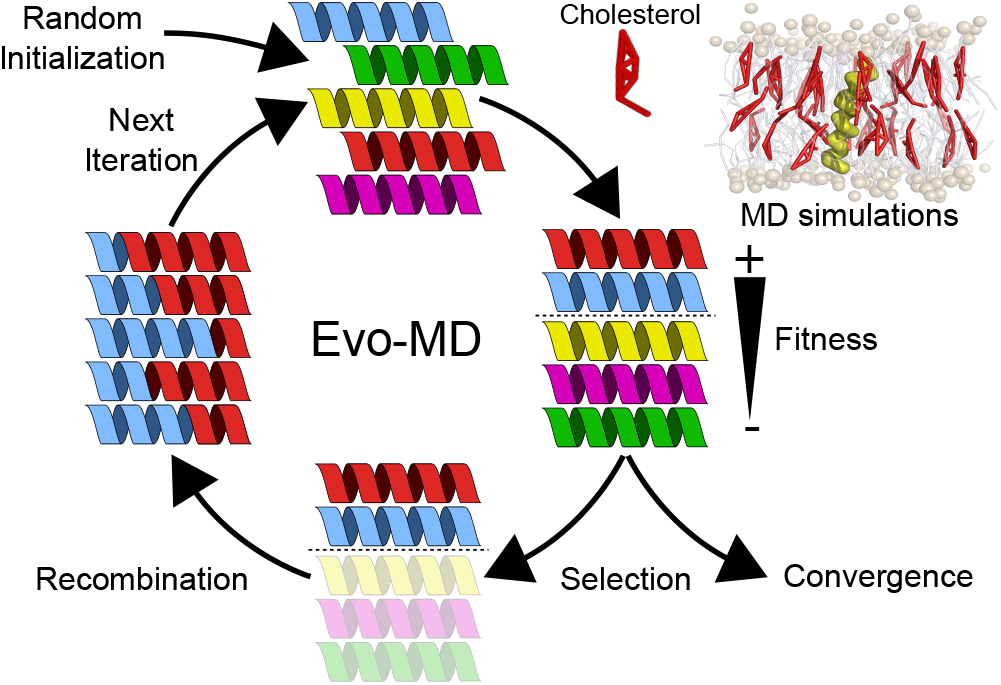
Illustration of the basic concept of evolutionary molecular dynamics (Evo-MD). Random peptide sequences self-evolve into optimal cholesterol attracting transmembrane domains in the course of evolution. Generated peptides are iteratively ranked upon increasing fitness, as determined via ensemble averaging within molecular dynamics simulations.

In this work, we apply Evo-MD to explore the evolutionary landscape of cholesterol attraction for a transmembrane domain sequence with a *fixed* length of 20 amino acids. We illustrate that cholesterol attraction is mediated by enforcing snorkeling of lipid head groups in the direct vicinity of the TMD, which energetically favors cholesterol over POPC lipids. In native membrane proteins, the presence of additional protein-protein interactions likely poses a far larger evolutionary constraint on sequence than protein-lipid interactions. Our observation of a single, well-defined minimum within sequence space suggests that cholesterol attraction in native TMDs is only sub-optimal, enabling cholesterol recognition motifs to be of a highly variable nature.

## Results

### Cholesterol sensing features a strong evolutionary conservation

Artificial evolution is simulated in a system consisting of a 30% cholesterol and 70% 1-palmitoyl-2-oleoyl-glycero-3phosphocholine (POPC) membrane containing a single, 20 amino acid long peptide sequence positioned transversely through the membrane (see Fig. 2A,B). Owing to the symmetry of the here-studied bilayer, generated sequences are mirror symmetric, i.e. only the first ten amino acids are independently chosen. Evolution is directed towards peptide sequences that increase the local density of cholesterol, visualized by the percentage cholesterol content of the membrane within a certain range from the peptide (see Fig. 2C). In practice, this is obtained by maximizing the ensemble-averaged non-bonded interaction energy between the peptide and cholesterol, i.e. this defines the *fitness*, in the course of sequence evolution.

**Fig. 2.**
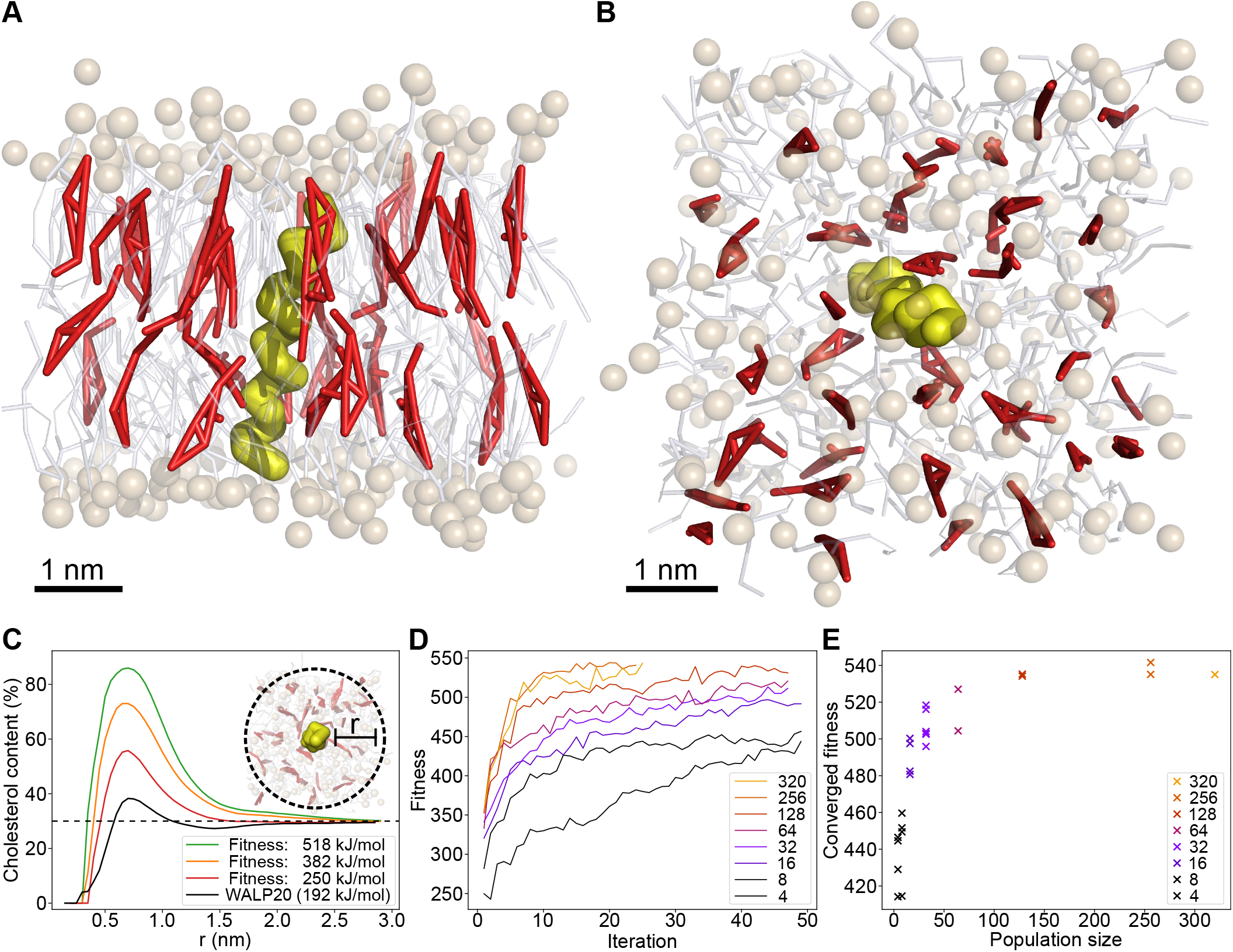
Evolutionary molecular dynamics simulations of a cholesterol attracting transmembrane protein. (A, B): Snapshots of a transmembrane protein (yellow) embedded within a POPC (white/brown) membrane containing 30% cholesterol (red). (C): Ratio of the cholesterol content in a local radius around the protein (see methods). An increase in fitness correlates to an increase in local cholesterol. The baseline cholesterol concentration (30%) is indicated by the dashed line. (D): Fitness development during protein evolution, shown for various population sizes. The fitness is expressed in terms of peptide-cholesterol interaction energy (Lennard-Jones energy). Fitness increases with GA iterations. Size of the population affects the height of the fitness plateau. (E): The GA converges to different fitness values, depending on the size of the populations. Eventually, evolution converges to an optimal solution for population sizes greater than 128 individuals.

Despite starting from *random* peptide sequences, the observed evolution eventually converges to an optimum, as is evident by a plateau in the fitness values (see Fig. 2D). Convergence of genetic algorithms depends on a variety of factors, most notably the size of the population—which directly relates to the area of the search space that is sampled each iteration—and the number of iterations that are performed.

Either parameter requires some minimum value for convergence to occur. The population size should be large enough (in combination with mutation rate and other diversifying factors) to prevent premature convergence to sub-optimal solutions, and, with evolution proceeding between iterations, a certain minimum number of iterations is necessary. Ideally, both parameters are chosen as large as possible.

To assess whether the convergence of evolution is either suboptimal (i.e, a local solution) or optimal (i.e., a global solution), we conducted a set of evolutionary runs with population sizes ranging from 4 to 320 individuals until no further convergence of fitness was observed. Figure 2D shows how the fitness of the best performing sequences changes with each generation. As expected, increasing population size increases the optimum fitness, as is evident from a higher plateau value reached after convergence of fitness (see Fig. 2E). This increase in optimal fitness leveled off once the population size began exceeding 128 individuals, which we took as the baseline population size for GA convergence. Data from GA runs containing 128+ individuals and at least 40 generations was used for sequence analysis.

Accompanying the convergence in fitness with respect to population size, we observed a similarity in sequences being produced by distinct GA runs. While GA runs with lower population sizes (*<* 64) did eventually converge to some fitness value, comparison between these distinct GA runs revealed a large diversity in the respective sequences, indicating that the algorithms converged to local optima in the solution space. This diversity in sequence decreases as population size increases, with very similar sequences being obtained as population sizes increase to 128 individuals and above. Furthermore, at such population sizes, starting the evolution from different initial populations consisting of randomly generated sequences yields a consistent result. On these grounds, we can conclude that the GA successfully converges to a global optimum.

To gain detailed insights into the resolved evolutionary landscape, high-fitness sequences from all GA runs with populations of 128+ individuals were combined to generate a sequence logo of the sampled sequence space (see Fig. 3A). Sequence logos express the degree of amino acid conservation at each position within the sequence in terms of the concomitant Shannon entropy (bits) by scaling the character height of the corresponding amino acid. Randomly occurring amino acids at a certain position contain no information, corresponding to a small letter, whereas a more frequently occurring amino acid encodes information, corresponding to a larger letter.

**Fig. 3.**
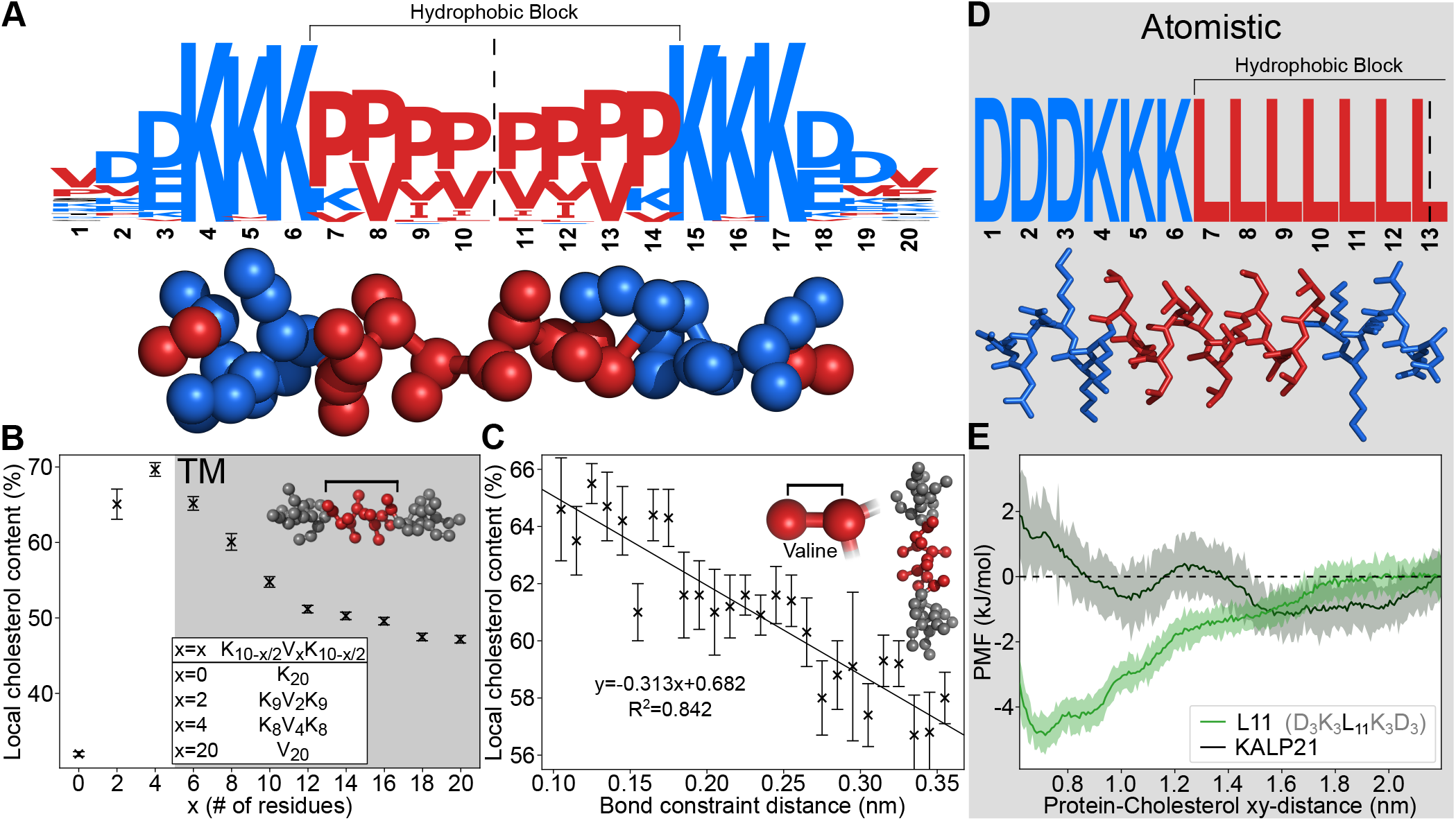
Sequence and chemical features of the optimal cholesterol attractor. (A): Sequence logo computed from all high-fitness (*≥* 500 kJ/mol) peptide sequences reveals a highly conserved hydrophobic pattern (red = hydrophobic; blue = hydrophilic). Owing to the symmetry of the here-used bilayer, sequences are mirrored around the center as indicated by the dashed line. (B): Short, hydrophobic blocks result in high local (1.0 nm radius) cholesterol composition of the membrane. Sequences adhere to the following motif: K_(10−*x*/2)_–V_*x*_–K_(10−*x*/2)_ (x=0, 2, 4 etc.). The gray region indicates the regime where a transmembrane (TM) topology is thermodynamically stable (see Fig. S10). Bars represent the standard error of the mean. (C): The cross-sectional thickness of proteins markedly affects the local cholesterol content (defined at a 1.0 nm radius) of the membrane surrounding the protein. Force field was adapted to vary the distance between valine backbone and side chain beads in a dummy peptide (sequence: K_6_V_8_K_6_). Bars represent the standard error of the mean. (D): A rationally designed motif (L11) based on the CG optimal cholesterol attractor. The sequence retains both the conserved poly-lysine patches, and the short hydrophobic section. The corresponding atomistic structure is shown below. (E): Free energy profiles over the peptide-cholesterol distance are computed in all-atom simulations for the rationally designed motif D_3_K_3_L_11_K_3_D_3_ (L11), and the stereotypical transmembrane peptide GK_2_[LA]_7_LK_2_A (KALP21). KALP21 is characterized by a slender hydrophobic motif rich in leucines [LA]_7_L. Nevertheless, a pronounced cholesterol attraction is only observed for the designed motif L11.

Interestingly, the global solution converges to a distinct pattern featuring a *short* conserved hydrophobic block consisting of proline (P), valine (V), and occasionally (iso)leucine (I/L), located in the center of the peptide, embedded within two hydrophilic blocks consisting of three highly conserved positively charged lysines (K). The first and last three terminal residues are less conserved and predominantly feature negatively charged aspartic acids (D). Finally, it is important to emphasize that the here-resolved solution space is subject to a constraint in secondary structure, i.e. all sequences are assumed to be alpha-helical (5). We will extensively address the transferability of solution space within a later section of this work.

### A short hydrophobic block is crucial for maximal cholesterol attraction

We observe the emergence of a specific hydrophobic pattern in the high-fitness peptide sequences. Most notable is the high conservation of the lysine (K) residues at positions 4–6 (and 15–17), which is conserved early in the evolution of the sequence and is present in all high fitness sequences. Surprisingly, a strong competition between lysine and arginine (R) is not observed, in contrast to their hypothesized equivalent role within the CRAC definition(18).

This sharp positional convergence of lysine prompted us to investigate what role the length of the hydrophobic block plays in the cholesterol sensing ability of the sequence. We created dummy peptides according to the K_(10−*x*/2)_–V_*x*_–K_(10−*x*/2)_ motif with each peptide consisting of 20 amino acids in total. Here, valines form the hydrophobic block of the peptides, with lysines functioning as the hydrophilic edges. By varying the number of valines and lysines, we effectively vary the length of the hydrophobic block.

Interestingly, cholesterol affinity increases with decreasing hydrophobic block length, with an optimal effect at a specific length (K_8_V_4_K_8_), see figure 3B. This pattern seems to arise from a trade-off between short block length and transmembrane (meta)stability, with a further decrease in block length resulting in a sharp decline in functionality. Artificially restraining a transmembrane orientation/topology for these motifs (K_9_V_2_K_9_, and even K_20_) restores the functionality to levels similar to the best performer (K_8_V_4_K_8_) (see Fig. S9). This suggests that cholesterol attraction is predominantly mediated by the lysine residues, conform with their high conservation, and that their positioning, i.e. the conserved hydrophobic block length, ensures a transmembrane topology in the course of evolution. Owing to the membrane thickening effect of cholesterol (29), cholesterol-enriched phases such as the liquid ordered (Lo) phase generally favors TMDs characterized by a long rather than short hydrophobic length (5, 30–33). This suggest that optimal targeting of free membrane cholesterol and cholesterol enriched ordered phases are, in fact, mediated by distinct physicochemical driving forces.

### Cholesterol clustering favors a slender hydrophobic block

A common ground between the resolved optimal sequences is that they are constructed from small hydrophobic amino acids, containing only a single side chain bead, suggesting that the GA seeks to optimize for cholesterol attraction by creating a thin, hydrophobic part within the peptide. This preference for small hydrophobic amino acid is further emphasized by the increased frequency of proline and valine compared to leucine and isoleucine, with the only difference between these amino acids within the coarse-grained forcefield being the bond length between the side chain and backbone bead (34). Despite its simplicity, this coarse-grained forcefield is able to predict preferential cholesterol binding to CRAC motifs present in the transmembrane domains of serotonin1A receptor and the ErbB2 growth factor receptor (22, 35).

We further investigate the preference for a thin hydrophobic block by creating a series of dummy peptides with sequence K_6_V_8_K_6_, where we vary the bond length of the valine residues, allowing us to artificially change the thickness of the hydrophobic block. Indeed, we observe that cholesterol attraction increases even further when we artificially decrease the side-chain bond distance of the valine residues, see figure 3C. Likewise, when we increase the bond distance, cholesterol attraction reduces. This is in line with our previous findings, and it is consistent with the notion that reduction of TMD surface area increases cholesterol affinity (5).

### The resolved hydrophobic pattern attracts cholesterol in atomistic simulations

In this work, we resolved the essential physicochemical driving forces that underpin cholesterol recognition in transmembrane domains within homogeneous model membranes. The here-resolved chemical features of the optimal cholesterol attractor are subsequently translated into realistic peptide sequences by anticipating for the following three model approximations:

I. Since transmembrane domains are predominantly alpha-helical, an alpha-helical secondary structure restraint was imposed on generated sequences. While this assumption simplifies the search space by circumventing the problem of secondary structure prediction, it introduces a possibility for amino acids with large penalties to alpha-helical propensity (e.g. proline, valine)(36) to occur in the generated sequences, producing non-helical peptides in unrestrained simulations. In line with the here-resolved properties of cholesterol attractors, i.e. the presence of a short and thin hydrophobic block, we devised a more realistic sequence by replacing these amino acids with helix-favoring alternatives such as leucine.
II. Electrostatic interactions are underestimated in the coarse-grained simulations, enabling the formation of sequences with a high net charge. To obtain a sequence with net zero charge, we balance the conserved lysines patches by adding three aspartic acids (D) to both terminal ends.
III. We anticipate on the notion that the coarse-grained model—and MD simulations in general—underestimate the length regime where transmembrane domains become thermodynamically stable with respect to experimental conditions. Transmembrane partitioning of polyleucine helices in experiments only becomes favorable over surface partitioning at a length of 10 leucines, in contrast to their atomistic estimation of 7-8 leucines(37) and our course-grained estimation of 6 leucines (see Fig. S10).

Altogether, this leads to the more realistic sequence D_3_K_3_L_11_K_3_D_3_ (L11), which retains all the design features put forth by the GA. Indeed, performing free energy calculations in atomistic MD simulations (see *Methods*) confirmed that such a sequence exhibits a pronounced cholesterol attracting functionality, as shown in figure 3E, particularly when compared with the prototypical and somewhat similar model peptide KALP21 (sequence: GKK(LA)_7_LKKA). We thus observe that the encoded functionality persists between the different model resolutions. Since the coarse-grained model is parameterized on reproducing thermodynamic properties while only coarsely capturing structural features, the observed independence on model resolution therefore strongly suggests that the mechanism behind this attraction is of a thermodynamic nature. Moreover, the obtained free energy profile illustrates that cholesterol attraction occurs over rather large distances—up to 1.8 nm—suggesting that the attraction is membrane mediated, and thus resulting from an interplay between peptide and membrane.

### Cholesterol attraction via linear motifs in nature

An interesting question is to what extent the two observed main features for cholesterol attraction —namely attraction via (i) reduction of the TMD’s surface area (e.g., selecting amino acid such as Valine, Leucine etc.) (5), and (ii) hydrophobic mismatch (e.g., relative positioning of Lysines within the TMD)—are being expressed in nature. Noting that hydrophobic mismatch is also a known determinant in protein trafficking as well as sorting (38, 39), one would therefore intuitively expect a stronger limitation on the evolutionary expression of such a mechanism. To this aim, we analyzed the cholesterol-attracting behavior of transmembrane motifs isolated from 8370 native membrane proteins in the TmAlphaFold database for membrane proteins(40), using a convolutional neural network (CNN) trained on fitness labeled data produced by EVO-MD (see methods and *SI*). It is important to emphasize that data generated by the directed evolution of random sequences encodes information on all possible thermodynamic driving forces relevant for cholesterol attraction.

Interestingly, we observed a slight difference between single-pass (bitopic) and multi-pass (polytopic) transmembrane proteins, with respect to both the fitness (Fig. 4A) and length of the TMD (Fig. 4B). In single-pass TMDs, the distribution of TMD lengths favors smaller TMDs (*<* 22 amino acids) compared to the multi-pass distribution, which is more narrowly centered around 22 amino acids. Remarkably, the distribution of fitness scores shows a slight shift for single-pass TMDs towards increased cholesterol attraction. This difference indeed suggests an increased evolutionary pressure towards the expression of linear motifs in single-pass TMDs over multi-pass TMDs, as cholesterol attracting residues can be ‘smeared out’ over neighboring transmembrane domains in multi-pass proteins (41).

**Fig. 4.**
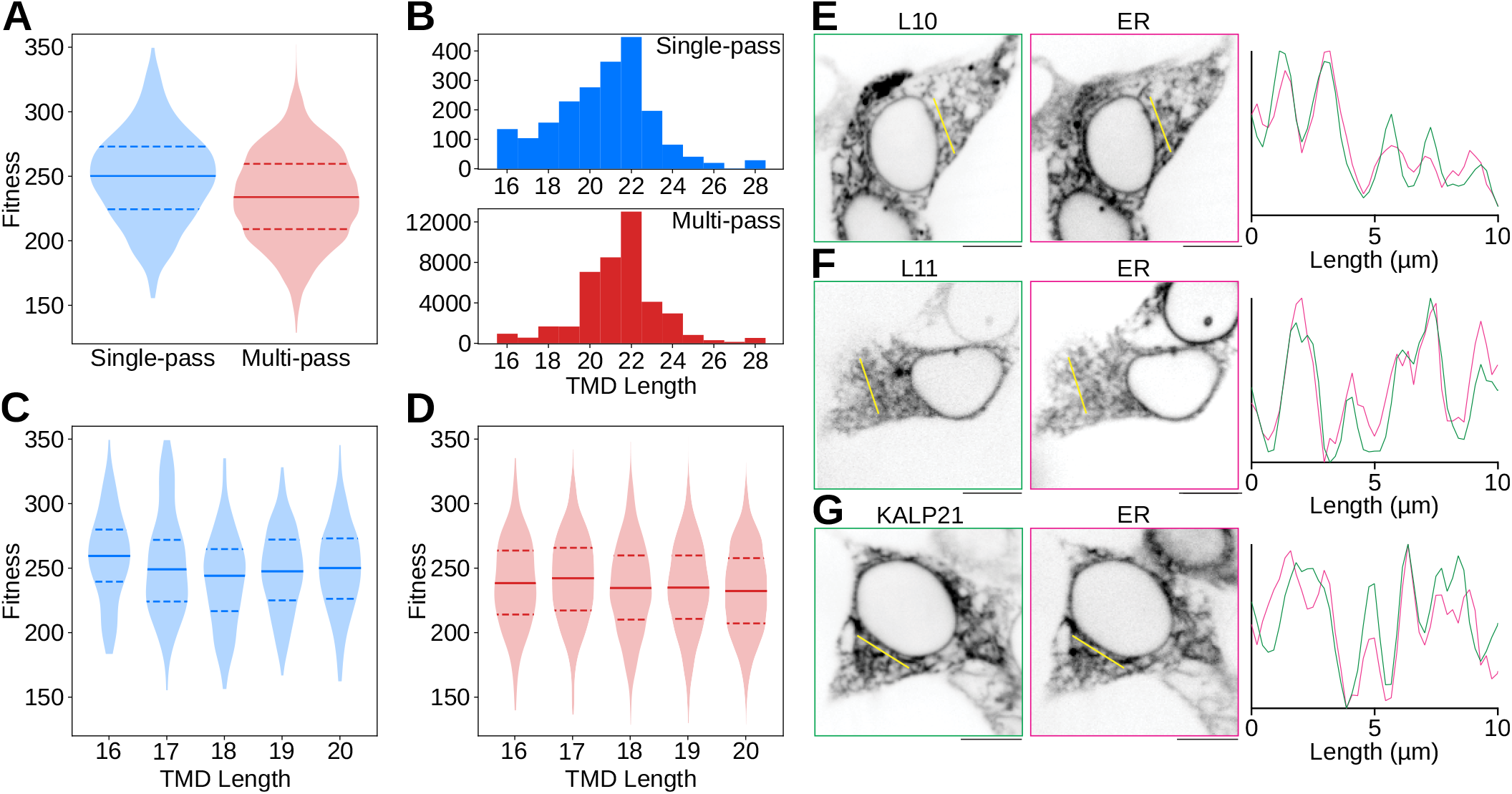
Existence and viability of mismatch-based attraction in nature. (A): Comparison of CNN-predicted fitness distributions between single-pass and multi-pass database TMDs. Markers indicate the interquartile range and the median of the data. (B): Frequency histograms of TMD length occurrences within the database. (C): Single-pass CNN-predicted fitness distributions with respect to TMD length. Markers indicate the interquartile range and the median of the data. (D): Multi-pass CNN-predicted fitness distributions with respect to TMD length. Markers indicate the interquartile range and the median of the data. (E-G): Fluorescence microscopy of transfected HEK cells, expressing L10 (E), L11 (F), and KALP21 (G); as well as fluorophore-tagged Sec61 to mark the ER. For each panel, a line profile was drawn (yellow), and the normalized fluorescence intensity profiles of peptide and ER are compared in the respective graphs. Scale bars and line profiles in all panels correspond to 10 μm.

However, no notable correlation was observed between predicted fitnesses and TMD length, in both the single-pass (Fig. 4C) and multi-pass datasets (Fig. 4D). This suggests that cholesterol attraction in nature is primarily based on amino acid composition (e.g., Lysines, Valines, Leucines etc.) rather than hydrophobic length. Thus, although physics-guided optimal cholesterol attraction is facilitated by the superposition of two distinct mechanisms, the hydrophobic mismatch mechanism is evidently being suppressed by evolutionary constraints in nature.

Even though hydrophobic mismatch driven attraction features optimal attraction at a hydrophobic length of 8 amino acids within our coarse-grained simulations, the shortest TMDs in the database still contained 16 amino acids (Fig. 4B). This implies that the optimal cholesterol attraction regime lies in fact far below the seemingly evolutionary accessible regime. To illustrate that extreme hydrophobic mismatch is physically achievable but not evolutionary viable, we performed experiments in live cells (HEK cells) expressing the short hydrophobic sequences D_3_K_3_L_10_K_3_D_3_ (L10) and D_3_K_3_L_11_K_3_D_3_ (L11), each with a fluorescent tag, as well as KALP21 (GK_2_[LA]_7_LK_2_A). KALP21 is a prototypical model peptide in membrane biophysical studies and has a (relatively short) hydrophobic length of 15 amino acids. Indeed, we observe that L10 (Fig. 4E), L11 (Fig. 4F), and KALP21 (Fig. 4G) can be successfully expressed by live cells and these transmembrane proteins all localize to the endoplasmic reticulum (ER), but not to other intracellular organelles (lysosomes/mitochondria) (Fig. S14), and not to the plasma membrane (Fig. S15). Nevertheless, the (over)expression of all three TMDs affected the transport of fatty acids and cholesterol from the ER membrane to the plasma membrane, as is demonstrated by altered localization of fat transporter and scavenger receptor CD36 (42, 43) within the plasma membrane (Fig S16), although we were unable to quantify individual differences therein.

Notably, the ER membrane is the thinnest membrane in live cells, and therefore shows the lowest energetic penalty for insertion of TMDs with negative hydrophobic mismatch (38, 44). However, the observation that even a stereotypical model protein such as KALP21 is restricted to the ER membrane, having only 1 amino acid less than the shortest native TMD within the TmAlphaFold database, underscores the dominant evolutionary barrier on exploiting the hydrophobic mismatch mechanism. Therefore, nature solely utilizes a amino acid composition mechanism, i.e. selecting aminoacids characteristic for the CRAC motif, to facilitate a thermodynamically sub-optimal but biologically sufficient attraction of cholesterol.

## Discussion

We have demonstrated the ability of Evo-MD to yield unique insights into the driving forces that underpin lipid recognition/binding in membrane proteins by resolving the actual thermodynamic optimum of cholesterol attraction. Physics-based inverse design of molecules is based on the principle that most if not all of the physical driving forces that govern functionality are inherently encoded within the complexity of independently parameterized classical molecular force fields. Our approach therefore substantially differs from popular data-driven quantitative structure-activity relationship (QSAR) based inverse design approaches, where optima in an abstract high-dimensional latent space are translated into corresponding chemical structures using machine learning-based variational encoders (26).

So far distinct motifs associated with cholesterol binding have been identified in native membrane proteins such as the CRAC motif and its inverse motif CARC(3). Arguably, one can include glycine zipper motifs such as found in the transmembrane domain of the amyloid precursor protein C99 (45), but where cholesterol binding is dominated by the additional presence of an N-terminal domain rather than the transmembrane domain itself (13). Surprisingly, we observe that optimal clustering of free membrane cholesterol to isolated transmembrane domains in fact prefers a short slender hydrophobic region flanked by lysine residues. This pattern evidently offers a pronounced net advantage for cholesterol binding in both coarse-grained and atomistic simulations.

The traditional lock and key hypothesis postulates that protein-ligand binding is mediated by matching of complementary molecular shapes. The optimal cholesterol attractor resolved in our Evo-MD simulations features a concave overall shape because of its voluminous hydrophilic ends and slender hydrophobic midsection. Interestingly, such a molecular shape is in principle compatible with the intrinsic cone-shape of cholesterol, i.e. small head group versus a voluminous hydrophobic moiety (46). Since both cholesterol and POPE lipids are cone-shaped molecules (46), and if binding is indeed effective molecular shape determined, then also POPE lipids are expected to bind the resolved motif. However, we do not observe an increased attraction for POPE lipids (see Fig. S12). As a matter of fact, a TMD optimized for the attraction of POPE lipids in fact features a highly contrasting structure (see Fig. 5C,D), i.e. a voluminous hydrophobic region comprised of tryptophans (W) with a slender intermediate hydrophilic region comprised of histidines (H). This suggests that the observed recognition of cholesterol molecules as well as POPE molecules is not based on effective molecular shape and is thus facilitated by a very different mechanism.

**Fig. 5.**
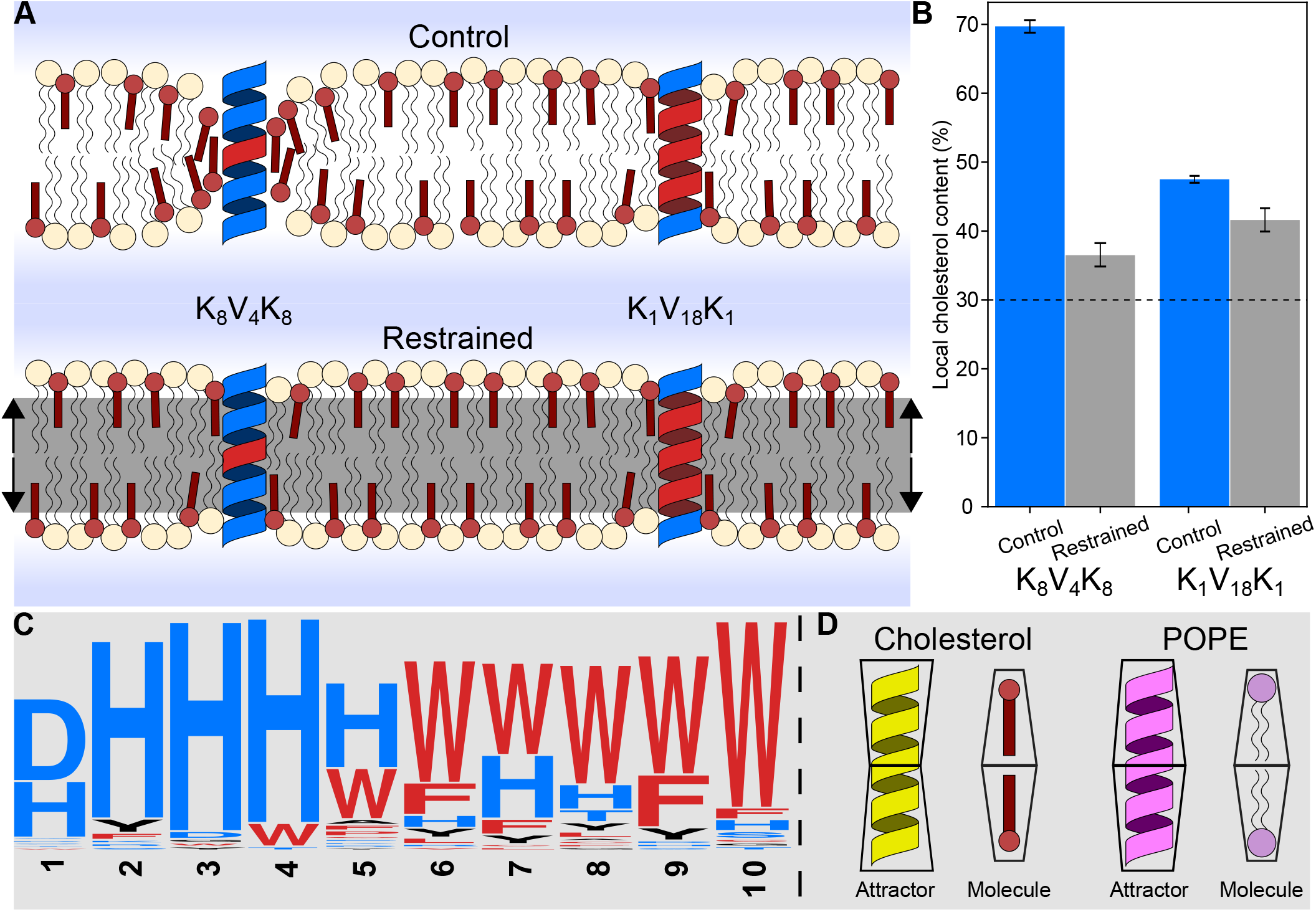
Mechanistic model for maximal cholesterol attraction. (A, Control): Cholesterol (red) has a lower energy penalty for membrane snorkeling compared to POPC (beige), allowing for more favorable shielding of deeply located lysine residues. High-affinity cholesterol attractors utilize this effect by increasing deep lysine interactions, leading to local accumulation of cholesterol molecules. (A, Restrained): Application of a force to lipid headgroups within a specific distance from the membrane center prevents cholesterol snorkeling/flip-flopping. (B): Removal of cholesterol snorkeling (Restrained) leads to a large drop in functionality for cholesterol attractors with short hydrophobic blocks (K_8_V_4_K_8_), while attractors with longer hydrophobic blocks (K_1_V_18_K_1_) are less affected. The dashed line indicates the average cholesterol content of the system (30%). Bars represent the standard error of the mean. (C): Sequence logo of a (mirrored) POPE attractor, computed from all high-fitness (*≥* 630 kJ/mol) peptide sequences, reveals a sequence distinct from the cholesterol attractor. (D): While cholesterol and POPE exhibit similarity in effective shape, corresponding attractor peptides vary notably in shape, suggesting a shape unspecific mechanism of attraction.

In contrast to POPC lipids, cholesterol faces a low free energy barrier to flip-flop between the two membrane leaflets. Consequently, the head group of cholesterol is relatively better at interacting with the deeply located lysine patch by snorkeling within the hydrophobic region of the membrane. Such positioning of cholesterol molecules shields the lysine patch from unfavorable interactions with the hydrophobic lipid tails (see Fig. 5A). Indeed, artificial restriction of lipid snorkeling confirms this hypothesis: high fitness sequences containing a short hydrophobic block—which would therefore highly depend on this snorkeling effect—show a large decrease in functionality, while ‘longer’ less optimal attractors—whose ability to attract cholesterol mostly relies on the slenderness of the hydrophobic section—remain relatively unaffected (see Fig. 5B). We therefore attribute the enhanced attraction of cholesterol to the enforced snorkeling of lipid head groups in the direct vicinity of the TMD. Within such a mechanism, a slender central hydrophobic TMD region comprised of, for example, leucines likely eases a closer binding of the cholesterol head groups to the lysine patch and thereby increases the associated enthalpic gain. Notably, also the optimal POPE attractor (see Fig. 5C) can be attributed to a differential difference in snorkeling between POPE and POPC lipids with the distinction that snorkeling now exploits a favorable enthalpic interaction between POPE head groups and the centrally located tryptophan region.

In essence, the thermodynamically optimal cholesterol attracting sequence can be regarded as a superposition of multiple CRAC motifs (3), excluding the aromatic residue tyrosine. Here it is important to emphasize that our sequence has evolved to maximize cholesterol clustering within the membrane, rather than specifically binding a single cholesterol molecule. Tyrosine can favorably bind the hydrophobic ring structure of a cholesterol molecule (3) and may thereby offer an advantage under the latter evolutionary pressure such as in the regulation of dimerization (3, 35). However, since tyrosine is a voluminous amino acid that is far less hydrophilic than lysine, stacking multiple tyrosines together with lysine seems counterproductive for maximizing cholesterol clustering. Notably, also the presence of multiple valines/leucines and a single lysine facilitates a pronounced cholesterol attraction even in the absence of tyrosine or a short hydrophobic block (Fig. 5B). This suggests that cholesterol binding can be variably encoded, in correspondence with findings in other studies (10, 15).

Surprising is the absence of arginine in cholesterol attractor sequences, with the CRAC motif suggesting a similar role for the lysine and arginine residues. Although substitution of lysine with arginine retains most of the functionality, a notable decrease in cholesterol attraction can be observed (see figure S11). This preference for lysine seems to stem from a larger difference in attraction/repulsion between cholesterol and POPC in the force field. While lysine and arginine have a charged residue end in common, the core of the residue differs, with arginine being more polar than lysine. The apolar lysine side chain provides a larger difference in attraction relative to arginine, with charged POPC headgroups being less attracted than the simply polar cholesterol headgroup. With the main feature of attraction satisfied by either residue (i.e., high hydrophilicity), we propose that this difference in side chain further discriminates between the two lipids. We hypothesize that such an effect is more pronounced in strong cholesterol clustering sequences, in contrast to the CRAC motif, due to the larger number of cholesterol molecules that are simultaneously attracted.

Our study focuses on cholesterol attraction within simple model membranes. However, verification with a coarse-grained model of the native epithelial membrane (47) illustrates that the here-resolved attraction mechanisms are universal and persist in more realistic membrane environments (see Fig. S8). In contrast, proteins targeting cholesterol-enriched liquid ordered phases (lipid rafts) are expected to prefer longer transmembrane domains (5, 30–33), likely to dampen hydrophobic mismatches between itself and the lipid rafts. The here-resolved motif is therefore not expected to optimally bind towards the interface of cholesterol-enriched liquid ordered domains (5, 48) (see Fig. S13). Nevertheless, the clustering of cholesterol is itself membrane phase independent and equally occurs when the resolved TMD is embedded within a liquid ordered DPPC:cholesterol mixture (see fig. S9).

Although the resolved hydrophobic pattern sequence adopts some of the basic features of the CRAC motif, the sequences of many native cholesterol-associated membrane proteins have evidently evolved on a respectable distance from the here-resolved thermodynamic attraction optimum. An obvious reason for this is that the true fitness function in nature will be subject to multiple distinct evolutionary pressures related to both protein trafficking, sorting and functionality, such as the potential interaction with raft domains and *especially* the interaction with other membrane proteins. Particularly, protein-protein interactions are highly specific and thus pose a far larger constraint on sequence variability than protein-lipid interactions. Although the here-resolved evolutionary landscape of cholesterol attraction features a well-defined and sharply peaked single optimum, the concomitant mechanism of attraction within lipid membranes is evidently not based on the precise matching of complementary molecular shapes, quite in contrast to typical ligand binding. Consequently, a large diversity of sub-optimal cholesterol attracting sequences can be achieved through evolution, explaining the puzzling variable nature of proposed CRAC motifs. Cholesterol binding to single-pass proteins is envisioned to mainly play a role in regulating dimerization (3, 35), mediating local membrane composition, and trafficking. Such functional binding seems indeed compatible with sequence unspecific thermodynamic attraction. However, we speculate that functional ligand-like binding of cholesterol to binding pockets formed between multipass transmembrane domains, dimers and higher oligomeric species will allow for a higher binding specificity and lower sequence variability since such a binding renders the matching of complementary molecular shapes more relevant over the here-observed thermodynamic mechanisms (7–10).

To summarize, we have demonstrated the ability of the Evo-MD approach to isolate the evolutionary fingerprints of protein-lipid interactions within membrane proteins. This unique property enables Evo-MD to yield valuable insights into *how* proteins may recognize and bind distinct membrane lipids or lipid soluble ligands such as hormones and vitamins within the highly crowded environment of lipid membranes. We therefore expect Evo-MD to yield novel insights into the mechanisms that underpin the molecular organization of biological membranes and protein trafficking. In particular, we have recently applied Evo-MD to the design of peptide motifs capable of selectively targeting membrane curvature(49), and are currently exploring the targeting of other characteristic membrane features such as lipid composition. Importantly, this will pave the road for the inverse design of peptide drugs and peptide based drug vehicles capable of selectively targeting the fluid membranes of viruses, microbes, and cancer cells; since their membrane leaflets are characterized by pronounced differences in curvature(50) and/or lipid composition(51). Notably, structure-based molecular design approaches are rendered ineffective beyond the level of targeting individual lipid species because of the diffusive, fluid nature of lipid membranes. Finally, as Evo-MD effectively extracts or mines implicit information from the underlying force field, integration of improved (coarse-grained) force fields—for example, the recent Martini 3 (52, 53) and the Spica force-field(54, 55)—in conjunction with the integration of groundbreaking protein structure prediction methodology—for example, the Alphafold 2 project (56)—could further facilitate these applications.

## Materials and Methods

### Software

Coarse-Grained simulations were performed with the Martini 2.2 CG force field using the GROMACS 2019.1 molecular dynamics package. EVO-MD is written in Python 3.6.8 and depends on the *NumPy* and *MPI for Python* packages for functionality. Peptide topologies are generated using *seq2itp* (34). Input parameters for the coarse-grained simulations are based on the Martini 2 ‘New-RF’ parameters (57), with exceptions detailed in the sections below.

### Implementation of simulated evolution: EVO-MD

EVO-MD was developed as a framework for the simulated evolution of MD simulation systems. Simulated evolution is a type of optimization problem involving the optimization of some property of the simulated system, by means of iteratively tuning a set of parameters. The performance (*fitness*) of such a parameter set is then measured by means of a fitness function, which generally consists of one or more MD simulations followed by an analysis step.

Using GAs, we can manage large, hyper-dimensional optimization problems through efficient exploration of the search space. Analogous to the method’s origin in genetics, we label each possible solution as a *chromosome*, which consists of a unique set of parameters encoded into a (bit)string sequence. The algorithm iteratively samples parts of the search space by forming a *population* of chromosomes and measuring their fitnesses. In line with evolution, individuals with high fitnesses are selected to *recombine* and form a new population. As the new population is based on a highest fitness subset of the previous population, it is assumed that the average fitness of the population increases each iteration. This process is visualized in figure 6.

**Fig. 6.**
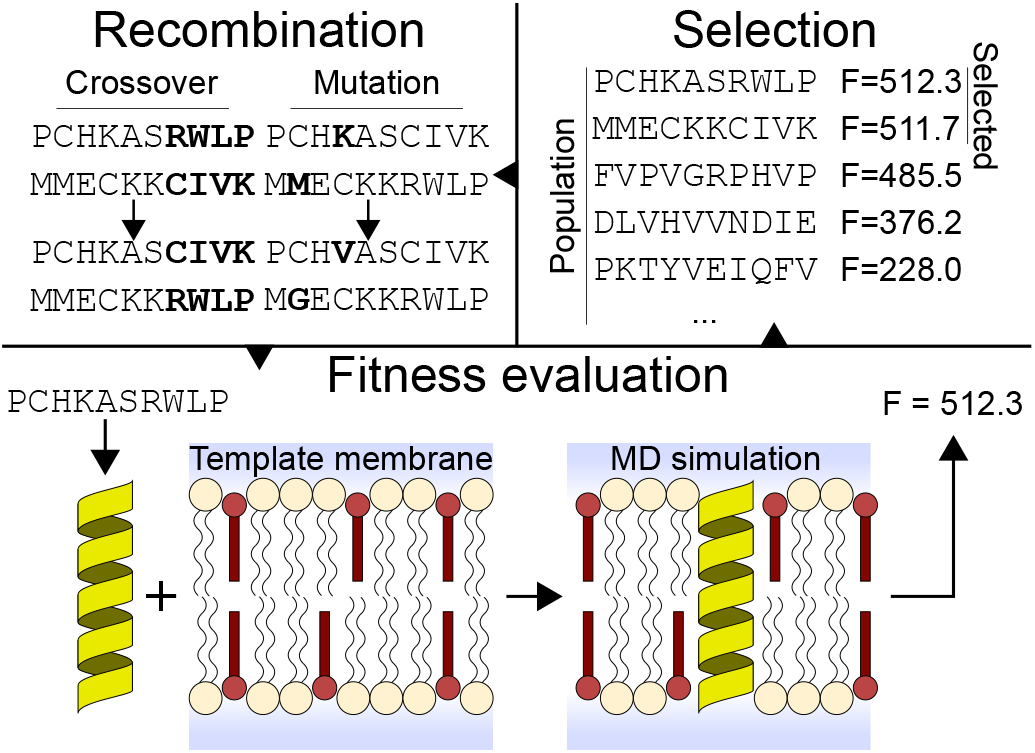
Graphical overview of EVO-MD. (Bottom): Peptide sequences are evaluated by means of MD simulation. A peptide structure (yellow) is generated from sequence and inserted into a POPC (beige) and cholesterol (red) bilayer membrane. The fitness is then computed from the resulting trajectory. (Top-right): Highest fitness sequences are selected from the evaluated population. (Top-left): Through recombination (involving crossover and mutation operations) of the selected sequences, a new population is generated.

Implementation of the cholesterol sensing project is summarized in figure S7. Each candidate peptide is encoded as a sequence of one-letter amino acid codes. For faster convergence, the sequence is mirrored to produce a palindromic sequence, effectively reducing the search space for a peptide 20 amino acids in length from 20^20^ to 20^10^ (assuming 20 amino acid types). The GA is initialized by generating a random population of *N*_*pop*_ sequences, after which each sequence is evaluated in parallel according to the fitness function.

The fitness function takes a sequence as argument and returns a single float value representing the sequence’s fitness. This function involves several simulation steps (*modules*): *generate_peptide, insert_peptide, production*, and *compute_fitness. Generate_peptide* generates a peptide structure and topology using the *seq2itp* tool (34), followed by energy minimization and peptide-membrane alignment. *Insert_peptide* combines the peptide structure with an existing equilibrated membrane structure containing 128 lipid molecules (90 POPC, 38 cholesterol) and 1598 Martini water beads, and places the peptide transversely through the membrane. Collisions between peptide and membrane structures are resolved by partially decoupling the non-bonded interactions—combined with soft-core potentials—and running a steepest descent algorithm. The *production* module adds ions to neutralize any net charge on the system, after which equilibration and production simulations are performed. The *compute_fitness* module then measures the ensemble-averaged short-ranged Lennard-Jones interactions between peptide and cholesterol (Coulomb interactions with cholesterol are absent within the CG model.) molecules from the simulation trajectory, which is returned as the fitness. Notable, such a fitness is the direct outcome of the competition between cholesterol and POPC lipids to interact with the peptide. Therefore, its value is directly proportional to the adopted cholesterol concentration and thus the *relative* binding free energy.

Once all sequences in the population have been evaluated, the algorithm proceeds by selecting the best *N* performers to serve as parents for the next population. A new sequence is generated by recombining two randomly selected sequences from the parent pool, which involves a *cross-over* operation and a *mutation* operation. During the cross-over operation, a random position is selected in the new sequence. The part to the left of that position is inherited from the first parent, while the rest of the sequence is inherited from the second parent. Afterwards, the mutation operation ensures that each position in the sequence has a 1*/len*(*sequence*) chance of being replaced with a random amino acid. New sequences are created in this manner until a new population of size *N*_*pop*_ is produced. This process of population fitness evaluation and recombination of the highest fitness candidates into a new population is then repeated until a desired number of iterations is achieved.

A *rerun* mechanism was implemented to account for possible undersampling during fitness evaluation. If a sequence reoccurs in a future generation, its fitness value will be determined as the weighted average of the current and all prior fitness evaluations. With the chance of sequence re-occurrence increasing as the algorithm converges, this mechanism serves to increase confidence in the final fitness value.

### Membrane setup

The membrane template structure consists of a 5.6×5.6×10 nm simulation box, containing a bilayer membrane in water solvent. The membrane consists of 90 POPC molecules and 38 cholesterol molecules. The solvent consists of 1598 Martini water beads.

### EVO-MD modules

#### Module: generate_peptide

As the seq2itp tool only produces topology files, a structure file for the peptide is generated by stacking hardcoded amino acid structures along the Z-axis and performing a 1.5 ps simulation at low time step (0.05 fs) using the GROMACS 2019.1 ‘sd’ stochastic dynamics integrator. This allows the hardcoded structure to slowly relax to a more reasonable conformation according to the generated topology.

#### Module: insert_peptide

*Insert_peptide* centers the peptide in the membrane box and merges the two structures together. A steepest descent, combined with a partial decoupling of the non-bonded interactions (*λ* = 0.75) and soft-core potentials, is then performed on the merged structure to remove collisions between the peptide and the membrane structures.

#### Module: production

A final steepest descent is performed without soft-core potentials. A short, 1.5 ps simulation is performed at low time step (0.05 fs) using the stochastic dynamics integrator to prevent blowing up of the system before the actual simulation is performed. The production simulation consists of a 500 ns NPT MD simulation with 30 fs time step, of which the first 50 ns are used for equilibration. Temperature is coupled to 300 K using velocity rescaling (*τ* = 1 ps with separate coupling groups for the membrane, peptide, and solvent), Pressure is coupled semi-isotropically to 1 bar using the Berendsen algorithm (*τ* = 8 ps), with compressibility set to 4.5 *·* 10^*−*5^ bar^*−*1^.

#### Module: compute_fitness

Evaluation of the sequence’s fitness is finalized by computation of a fitness value from the produced simulation trajectory. GROMACS’ *gmx energy* tool is used to extract the ensemble average of the non-bonded interaction energies from the production trajectory. The absolute value is then returned to the GA.

Quantification of sequence cholesterol clustering capability was performed by measuring the ratio of cholesterol molecules to membrane molecules within a cylinder of radius *r* centered on the peptide center-of-mass (COM). GROMACS’ *gmx rdf* tool was used to compute a cumulative number radial distribution function (*g*_*CN*_ (*r*)) for cholesterol COMs and POPC COMs, both with respect to the peptide COM. The final ratio figures are created by computing 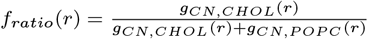.

Comparisons between multiple ratio figures were taken at a cylinder radius of 1.0 nm, chosen as a middle-ground between local-sampling (low *r*) and sufficient sampling (high *r*).

### GA parameters

Production runs of the GA were performed according to the parameters as described in Table 1. *Parents* indicates the size of the selection pool, from which parents were selected at random for the recombination step. *Iteration elites* describe the number of highest fitness sequences which pass unaltered into the next generation. *Rerun elites* keeps track of a list of sequences which have been evaluated more than once, and allows several highest fitness sequences to proceed to the next generation unaltered. The total number of elites is equal to the sum of iteration and rerun elites.

**Table 1.** Overview of GA run parameters.

| Population | of GA runs | Parents | Iteration elites | Rerun elites | Mutation frequency* |
| --- | --- | --- | --- | --- | --- |
| 4 | 4 | 2 | 1 | 1 | 1/20 |
| 8 | 4 | 2 | 1 | 1 | 1/20 |
| 16 | 4 | 4 | 1 | 1 | 1/20 |
| 32 | 4 | 8 | 2 | 2 | 1/20 |
| 64 | 2 | 16 | 2 | 2 | 1/20 |
| 128 | 2 | 16 | 2 | 2 | 1/20 |
| 256 | 2 | 16 | 2 | 2 | 1/20 |
| 320 | 1 | 16 | 2 | 2 | 1/20 |

\*per amino acid

### Free energy calculations in all-atom simulations

Simulations were performed using the GROMACS 2021.3 molecular dynamics package, with the Plumed 2.7.2 plugin. The KALP21 and L11 peptides were represented using AMBER99SB-ILDN(58), while POPC and cholesterol were represented with the Slipids forcefield(59, 60). For water molecules we used the TIP3P model. Simulations were performed in the NPT ensemble at 303.15 K, maintained with a Nose-Hoover thermostat. Pressure was kept at 1 bar using a semi-isotropic coupling scheme and a Parrinello-Rahman barostat. Long-range electrostatic interactions were calculated using the PME algorithm with a real-space cutoff of 1.4 nm. Van der Waals interactions were calculated with a 1.4 nm cutoff, and dispersion corrections for energy and pressure were applied. The leap-frog algorithm with a time step of 2 fs was used to integrate equations of motion. The LINCS algorithm was used to constrain hydrogen atom-containing bonds.

To determine the free energy profile for the cholesterol binding to KALP21 and L11, we used umbrella sampling (US). As the reaction coordinate, we used the in-plane center-of-mass distance (xy-distance) between the cholesterol ring system and all the peptide C*α* atoms located in the same membrane leaflet as the cholesterol molecule (residues 1–11 and 1–12 for KALP21 and L11, respectively). To describe the binding process, we sampled the 0.7–2.3 nm range of xy-distance using 9 evenly spaced US windows separated by 0.2 nm. In each window the reaction coordinate was subject to a harmonic bias potential with a spring constant of 250 kJ ·mol^*−*1^ · nm^*−*2^. For each window, 1.5 μs simulations were performed, and the the free energy profiles were calculated using the WHAM method. For each window, the first 400 ns of the trajectory were discarded as equilibration. The statistical uncertainties of the free energy were estimated using the Monte Carlo bootstrap method, taking into account autocorrelation times.

### Restraining lipid snorkeling/flip-flopping

To investigate the cholesterol snorkeling mechanism, removal of lipid snorkeling and flip-flopping was facilitated by application of an inverse flat-bottomed position restraint to the first beads of both POPC (NC3 bead) and cholesterol (ROH bead). The position restraint consists of a layer, parallel to the membrane and centered on the bilayer center. A harmonic force with force constant 1000 kJ ·mol^*−*1^ · nm^*−*2^, directed away from the bilayer center, is applied to affected beads that come within 2.0 nm (NC3) or 1.5 nm (ROH) of the center of the bilayer.

### Database analysis using a convolutional neural network

#### Convolutional neural network

The CNN architecture consisted of a one-hot encoding step, which is fed into 2 convolutional layers (128 nodes each) with max pooling, followed by 2 fully-connected dense layers (36 nodes each) and a single output neuron. The random dropout, which is applied before the output of the convolutional layers enters the dense layers, was set to 0.5 %. The model was trained in 16 epochs, with a batch size of 64 and a learning rate of 0.001.

#### Database analysis

Protein sequences and corresponding transmembrane predictions were downloaded from the TmAlphaFold Trans-membrane Protein Structure Database (https://tmalphafold.ttk.hu/downloads). From this database, *Homo sapiens* (UP000005640), *Mus musculus* (UP000000589), and *Rattus norvegicus* (UP000002494) were considered for analysis. We only included proteins that passed all 10 TM prediction quality flags (i.e. categorized as ‘excellent’), as described in (40). The resulting dataset contained 8370 protein entries in total, which was subsequently split in a single-pass dataset (2084 entries) and a multi-pass dataset (6286 entries, 42436 passes). We post-processed these datasets to produce sequences of 20 amino acids, as the CNN was trained on this type of data. TM sequences that exceeded 20 amino acids in length were removed, and TM sequences shorter than 20 amino acids were extended evenly along the edges using the corresponding non-TM amino acids from the protein sequence. We ended up with 902 single-pass sequences and 11954 multi-pass sequences, which we used for fitness prediction using the CNN, and subsequent analysis.

### In-vitro expression of short hydrophobic sequences

#### Cloning

Amino acid sequences for D_3_K_3_L_10_K_3_D_3_ (L10), D_3_K_3_L_11_K_3_D_3_ (L11), and GK_2_[LA]_7_LK_2_A (KALP21) were introduced into mScarlet-N1 and mEmerald-N1 by Gibson assembly. The final constructs were all confirmed by sequencing.

#### Cell-based experiments

HEK cells were transfected by Lipofectamine 2000 (Thermo Fisher Scientific) following manufacturer’s instructions. Briefly, 3 ml of Lipofectamine 2000 was mixed with max 2 mg of total DNA (in equimolar ratio) in 200 ml OptiMEM (Gibco). Transfection mix was incubated for 30 min at room temperature, then was added to the cells. Cells were transfected and incubated overnight (37 °C and 5% CO2). The day after medium was fully replaced with fresh supplemented DMEM.

Prior to imaging the transfected cells were incubated with Wheat Germ Agglutinin, CF®405S Conjugate (WGA405, 1:20, stock 2 mg/ml) for 10 min. During acquisition the laser power was kept constant. Exposure time 200 ms and Piezo stage z-motor was used to collect z-stacks.

For visualizing the intracellular organelles, the transfected cells were incubated with LysoTracker™ Deep Red, (Thermo Fisher Scientific) and MitoTracker™ Deep Red FM, (Thermo Fisher Scientific) for visualization of lysosomes and mitochondria, respectively. Fluorescent dyes were diluted 1:1,000 in pre-warmed imaging solution and added to HEK cells 10 min before imaging.

#### Image analysis

Images were acquired using Acquisition software NIS Elements 5.21.02 and analyzed with ImageJ (NIH). Freehand selection tool in Fiji was used to select a region of interest (ROI) of 3 pixel width following the fluorescence signal of WGA405 (i.e. 405-channel) as reference for plasma membrane from the medial cell plane. Each ROI was assessed for the signal intensity in the 561-channel (for mCherry-CD36) and the mean fluorescent intensity was measured from the ROI for calculating Intensity/length (mm). GraphPad Prism 9 was used to plot the graphs (each value is shown as the average ± standard error of the mean). Statistical test performed was unpaired t-Test (“P value”<0.0001”, P value summary ****).

## ACKNOWLEDGMENTS

The authors gratefully acknowledge the Gauss Centre for Supercomputing e.V. (www.gauss-centre.eu) for funding this project by providing computing time through the John von Neumann Institute for Computing (NIC) on the GCS Supercomputer JUWELS at Jülich Supercomputing Centre (JSC).

## Notes

### Competing Interest Statement

The authors have declared no competing interest.

### Summary of Updates

Included acknowledgements.

## References

1. Levental I, Veatch SL (2016) The continuing mystery of lipid rafts. J. Mol. Biol. 428(24):4749–4764.

2. Midzak A, Papadopoulos V (2014) Binding domain-driven intracellular trafficking of sterols for synthesis of steroid hormones, bile acids and oxysterols. Traffic 15(9):895–914.

3. Fantini J, Barrantes FJ (2013) How cholesterol interacts with membrane proteins: an exploration of cholesterol-binding sites including CRAC, CARC, and tilted domains. Front. Physiol. 4.

4. Scala CD, et al. (2017) Relevance of CARC and CRAC cholesterol-recognition motifs in the nicotinic acetylcholine receptor and other membrane-bound receptors. Curr. Top. Membr. 80:3–23.

5. Lorent JH, et al. (2017) Structural determinants and functional consequences of protein affinity for membrane rafts. Nat. Commun. 8(1):1219.

6. Fatakia SN, Sarkar P, Chattopadhyay A (2019) A collage of cholesterol interaction motifs in the serotonin1a receptor: An evolutionary implication for differential cholesterol interaction. Chem. Phys. Lipids 221:184–192.

7. Bukiya AN, Dopico AM (2017) Common structural features of cholesterol binding sites in crystallized soluble proteins. J. Lipid Res. 58(6):1044–1054.

8. Wang C, Ralko A, Ren Z, Rosenhouse-Dantsker A, Yang X (2019) Modes of cholesterol binding in membrane proteins: a joint analysis of 73 crystal structures. Adv. Exp. Med. Biol. pp. 67–86.

9. Dubey V, Bozorg B, Wüstner D, Khandelia H (2020) Cholesterol binding to the sterol-sensing region of niemann pick c1 protein confines dynamics of its n-terminal domain. PLoS Comp. Biol. 16(10):e1007554.

10. Marlow B, Kuenze G, Li B, Sanders CR, Meiler J (2021) Structural determinants of cholesterol recognition in helical integral membrane proteins. Biophysical Journal 120(9):1592–1604.

11. Luo J, Yang H, Song BL (2019) Mechanisms and regulation of cholesterol homeostasis. Nat. Rev. Mol. Cell Biol. 21(4):225–245.

12. Li H, x. Yao Z, Degenhardt B, Teper G, Papadopoulos V (2001) Cholesterol binding at the cholesterol recognition/ interaction amino acid consensus (CRAC) of the peripheral-type benzodiazepine receptor and inhibition of steroidogenesis by an HIV TAT-CRAC peptide. Proc. Natl. Acad. Sci. USA 98(3):1267–1272.

13. Nierzwicki Ł, Czub J (2015) Specific binding of cholesterol to the amyloid precursor protein: structure of the complex and driving forces characterized in molecular detail. J. Phys. Chem. Lett. 6(5):784–790.

14. Koufos E, Chang EH, Rasti ES, Krueger E, Brown AC (2016) Use of a cholesterol recognition amino acid consensus peptide to inhibit binding of a bacterial toxin to cholesterol. Biochemistry 55(34):4787–4797.

15. Elkins MR, et al. (2017) Cholesterol-binding site of the influenza m2 protein in lipid bilayers from solid-state NMR. Proc. Natl. Acad. Sci. USA 114(49):12946–12951.

16. Castellano BM, et al. (2017) Lysosomal cholesterol activates mTORC1 via an SLC38a9–niemann-pick c1 signaling complex. Science 355(6331):1306–1311.

17. Epand RM (2006) Cholesterol and the interaction of proteins with membrane domains. Prog. Lipid Res. 45(4):279–294.

18. Fantini J, Epand RM, Barrantes FJ (2019) Cholesterol-recognition motifs in membrane proteins. Adv. Exp. Med. Biol. 1135:3–25.

19. Rosenhouse-Dantsker A, Noskov S, Durdagi S, Logothetis DE, Levitan I (2013) Identification of novel cholesterol-binding regions in kir2 channels. J. Biol. Chem. 288(43):31154–31164.

20. Singh AK, et al. (2012) Multiple cholesterol recognition/interaction amino acid consensus (CRAC) motifs in cytosolic c tail of slo1 subunit determine cholesterol sensitivity of ca2- and voltage-gated k(BK) channels. J. Biol. Chem. 287(24):20509–20521.

21. Jafurulla M, Tiwari S, Chattopadhyay A (2011) Identification of cholesterol recognition amino acid consensus (CRAC) motif in g-protein coupled receptors. Biochem. Biophys. Res. Commun. 404(1):569–573.

22. Sengupta D, Chattopadhyay A (2012) Identification of cholesterol binding sites in the serotonin1a receptor. J. Phys. Chem. B 116(43):12991–12996.

23. Hedger G, et al. (2019) Cholesterol interaction sites on the transmembrane domain of the hedgehog signal transducer and class f g protein-coupled receptor smoothened. Structure 27(3):549–559.e2.

24. Sejdiu BI, Tieleman DP (2020) Lipid-protein interactions are a unique property and defining feature of g protein-coupled receptors. Biophys. J. 118(8):1887–1900.

25. Lutz S (2010) Beyond directed evolution—semi-rational protein engineering and design. Curr. Opin. Biotechnol. 21(6):734–743.

26. Sanchez-Lengeling B, Aspuru-Guzik A (2018) Inverse molecular design using machine learning: Generative models for matter engineering. Science 361(6400):360–365.

27. Kaelbling LP, Littman ML, Moore AW (1996) Reinforcement learning: A survey. Journal of Artificial Intelligence Research 4:237–285.

28. Sloss AN, Gustafson S (2019) 2019 evolutionary algorithms review.

29. Risselada HJ, Marrink SJ (2008) The molecular face of lipid rafts in model membranes. Proc. Natl. Acad. Sci. USA 105(45):17367–17372.

30. Schafer LV, et al. (2011) Lipid packing drives the segregation of transmembrane helices into disordered lipid domains in model membranes. Proc. Natl. Acad. Sci. USA 108(4):1343–1348.

31. Kaiser HJ, et al. (2011) Lateral sorting in model membranes by cholesterol-mediated hydrophobic matching. Proc. Natl. Acad. Sci. USA 108(40):16628–16633.

32. Milovanovic D, et al. (2015) Hydrophobic mismatch sorts SNARE proteins into distinct membrane domains. Nat. Commun. 6(1).

33. Chakraborty S, et al. (2020) How cholesterol stiffens unsaturated lipid membranes. Proc. Natl. Acad. Sci. USA 117(36):21896–21905.

34. Monticelli L, et al. (2008) The martini coarse-grained force field: Extension to proteins. Journal of Chemical Theory and Computation 4(5):819–834.

35. Pawar AB, Sengupta D (2021) Role of cholesterol in transmembrane dimerization of the erbb2 growth factor receptor. The Journal of Membrane Biology 254:301–310.

36. Nick Pace C, Martin Scholtz J (1998) A helix propensity scale based on experimental studies of peptides and proteins. Biophysical Journal 75(1):422–427.

37. Ulmschneider JP, Smith JC, White SH, Ulmschneider MB (2011) In silico partitioning and transmembrane insertion of hydrophobic peptides under equilibrium conditions. Journal of the American Chemical Society 133(39):15487–15495.

38. Mitra K, Ubarretxena-Belandia I, Taguchi T, Warren G, Engelman DM (2004) Modulation of the bilayer thickness of exocytic pathway membranes by membrane proteins rather than cholesterol. Proc. Natl. Acad. Sci. U.S.A 101(12):4083–4088.

39. Milovanovic D, et al. (2015) Hydrophobic mismatch sorts snare proteins into distinct membrane domains. Nat. Commun. 6(1):1–10.

40. Dobson L, et al. (2022) TmAlphaFold database: membrane localization and evaluation of AlphaFold2 predicted alpha-helical transmembrane protein structures. Nucleic Acids Research. gkac928.

41. Najbauer EE, et al. (2021) Structure, gating and interactions of the voltage-dependent anion channel. EU. Biophys. J. 50(2):159–172.

42. Wang J, et al. (2019) DHHC4 and DHHC5 facilitate fatty acid uptake by palmitoylating and targeting CD36 to the plasma membrane. Cell Rep 26(1):209–221.e5.

43. Hao JW, et al. (2020) CD36 facilitates fatty acid uptake by dynamic palmitoylation-regulated endocytosis. Nat Commun 11(1):4765.

44. Singh S, Mittal A (2016) Transmembrane domain lengths serve as signatures of organismal complexity and viral transport mechanisms. sci rep 6: 22352.

45. Barrett PJ, et al. (2012) The amyloid precursor protein has a flexible transmembrane domain and binds cholesterol. Science 336(6085):1168–1171.

46. Kollmitzer B, Heftberger P, Rappolt M, Pabst G (2013) Monolayer spontaneous curvature of raft-forming membrane lipids. Soft matter 9(45):10877–10884.

47. Wilson KA, et al. (2021) The role of plasmalogens, forssman lipids, and sphingolipid hydroxylation in modulating the biophysical properties of the epithelial plasma membrane. The Journal of Chemical Physics 154(9):095101.

48. Risselada HJ, Marrink SJ (2008) The molecular face of lipid rafts in model membranes. Proceedings of the National Academy of Sciences 105(45):17367–17372.

49. van Hilten N, Methorst J, Verwei N, Risselada HJ (2023) Physics-based generative model of curvature sensing peptides; distinguishing sensors from binders. bioRxiv.

50. Yoon BK, Jeon WY, Sut TN, Cho NJ, Jackman JA (2021) Stopping membrane-enveloped viruses with nanotechnology strategies: Toward antiviral drug development and pandemic preparedness. ACS Nano 15(1):125–148.

51. Rivel T, Ramseyer C, Yesylevskyy S (2019) The asymmetry of plasma membranes and their cholesterol content influence the uptake of cisplatin. Scientific Reports 9(1).

52. Souza PC, et al. (2021) Martini 3: a general purpose force field for coarse-grained molecular dynamics. Nat. Method. pp. 1–7.

53. Risselada HJ (2021) Martini 3: a coarse-grained force field with an eye for atomic detail. Nat. Methods. 18(4):342–343.

54. Shinoda W, DeVane R, Klein ML (2007) Multi-property fitting and parameterization of a coarse grained model for aqueous surfactants. Molecular Simulation 33(1-2):27–36.

55. Seo S, Shinoda W (2019) Spica force field for lipid membranes: Domain formation induced by cholesterol. J. Chem. Theory Comput. 15(1):762–774. PMID: 30514078.

56. Senior AW, et al. (2020) Improved protein structure prediction using potentials from deep learning. Nature 577(7792):706–710.

57. de Jong DH, Baoukina S, Ingólfsson HI, Marrink SJ (2016) Martini straight: Boosting performance using a shorter cutoff and gpus. Computer Physics Communications 199:1–7.

58. Lindorff-Larsen K, et al. (2010) Improved side-chain torsion potentials for the amber ff99sb protein force field. Proteins: Structure, Function, and Bioinformatics 78(8):1950–1958.

59. Jämbeck JPM, Lyubartsev AP (2012) An extension and further validation of an all-atomistic force field for biological membranes. Journal of Chemical Theory and Computation 8(8):2938–2948.

60. Jämbeck Jpm, Lyubartsev AP (2012) Another piece of the membrane puzzle: Extending slipids further. Journal of Chemical Theory and Computation 9(1):774–784.

