## Supplementary Information for "Physics-based inverse design of cholesterol attracting transmembrane helices reveals a paradoxical role of hydrophobic length"

### Methods.

***Native epithelial membrane.*** A cholesterol attractor ($D_3K_3L_8K_3D_3$) was inserted into a native epithelial membrane model ((47), "Membrane 1") and a 2 $\mu$s NPT MD simulation is performed with a 20 fs time step, of which the first 500 ns were used for equilibration purposes. Temperature was coupled to 310 K using velocity rescaling ($\tau = 1$ ps, with separate coupling groups for the membrane, peptide, and solvent). Pressure was coupled semi-isotropically to 1 bar using the Parrinello-Rahman barostat ($\tau = 12$ ps), with compressibility set to $3.0 * 10^{-4}$ bar$^{-1}$.

Cholesterol content was computed from the ratio of cholesterol molecules to membrane molecules within a cylinder of radios $r$ centered on the peptide center-of-mass (COM) (i.e. $f_{ratio}(r) = \frac{g_{CN,CHOL}(r)}{g_{CN,Lipids}(r)}$).

***Restraining peptide transmembrane position.*** Several peptides with unfavorable transmembrane affinity have been restrained to better investigate the underlying effect of hydrophobic block length on cholesterol attraction. After a peptide structure was inserted into the template membrane, a short 5 ns equilibration step was performed to position the peptide. The first and last backbone beads of the peptide were then restrained using a flat-bottom potential to 0.25 nm thick layers parallel to the membrane, relative to their equilibrated positions. To prevent the membrane from adjusting to the peptides enforced position, all lipid beads were restrained to a 5.52 nm thick layer centered and parallel to the membrane.

***Free-energy calculations.*** To determine the transmembrane stability of peptides, we computed the free energy of insertion ($\Delta G_{insertion}$) using the thermodynamic integration (TI) method. For both flat and TM configurations, coulomb and Van der Waals interactions were decoupled separately in steps of $\lambda = 0.05$. The free energy then becomes $\Delta G_{insertion} = \Delta G_{flat,coul} + \Delta G_{flat,vdw} - \Delta G_{TM,vdw} - \Delta G_{TM,coul}$.

**Movies.** Directed evolution leads to a convergence in sequence. For each iteration, a sequence logo was generated using all sequences from GA runs with population sizes $\geq 128$.

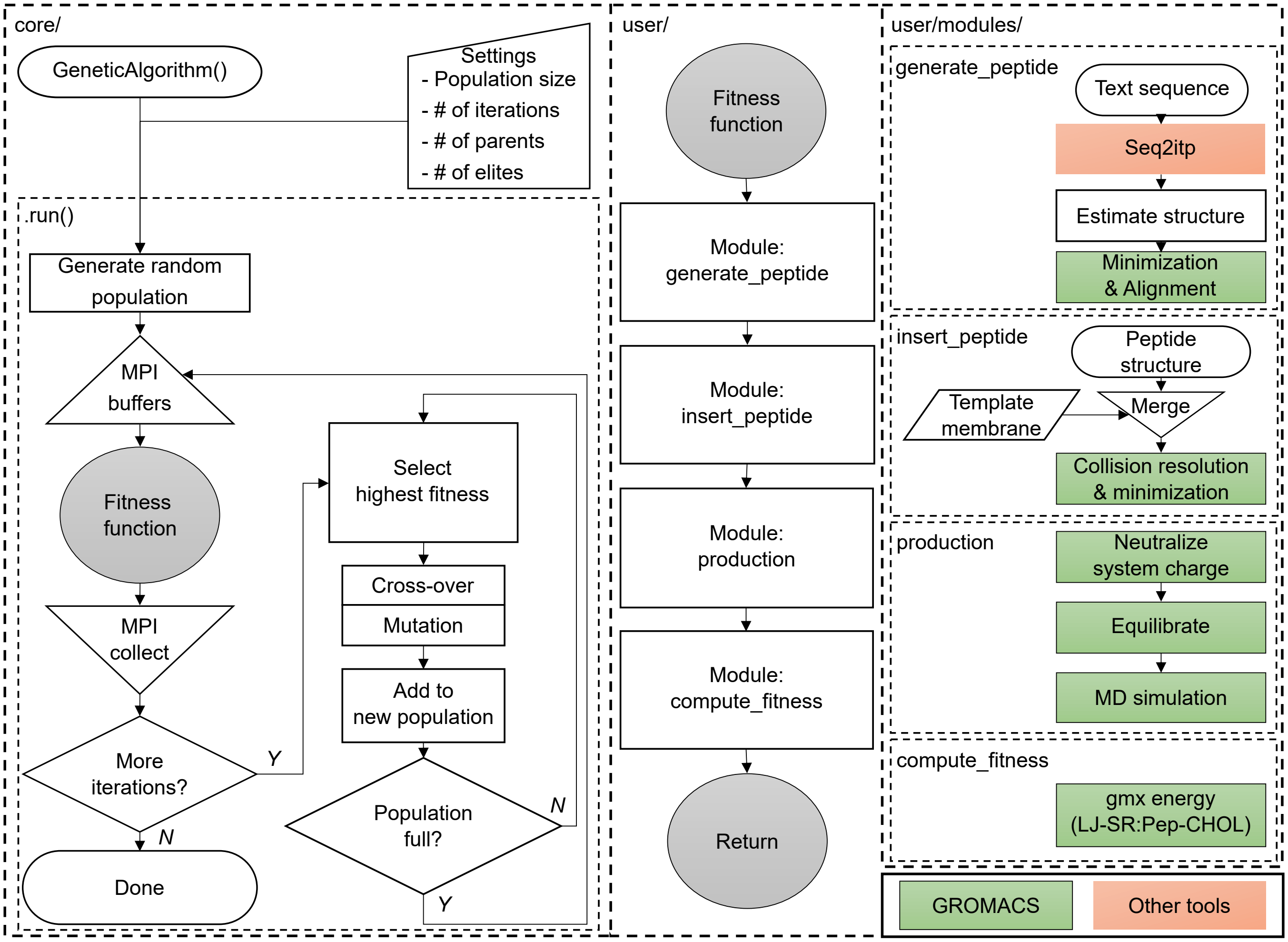


**Fig. S7.** Flowchart of EVO-MDimplementation. (core/) Genetic algorithm core of EVO-MD. An instance of the GeneticAlgorithm class is made and initialized with run parameters (e.g. population size, sequence length, etc...). The population is then initialized with random sequences and evaluated over all available MPI ranks using the fitness function. Best performing sequences are selected based on fitness, and a new population is formed using cross-over and mutation operators. (user/ and user/modules/) Cholesterol attractor implementation into EVO-MD. Fitness function receives a peptide sequence from the GA from which a simulation system is generated. Using the resulting simulation trajectory, a fitness value is computed based on non-bonded interaction energies between cholesterols and peptide, which is returned to the GA.

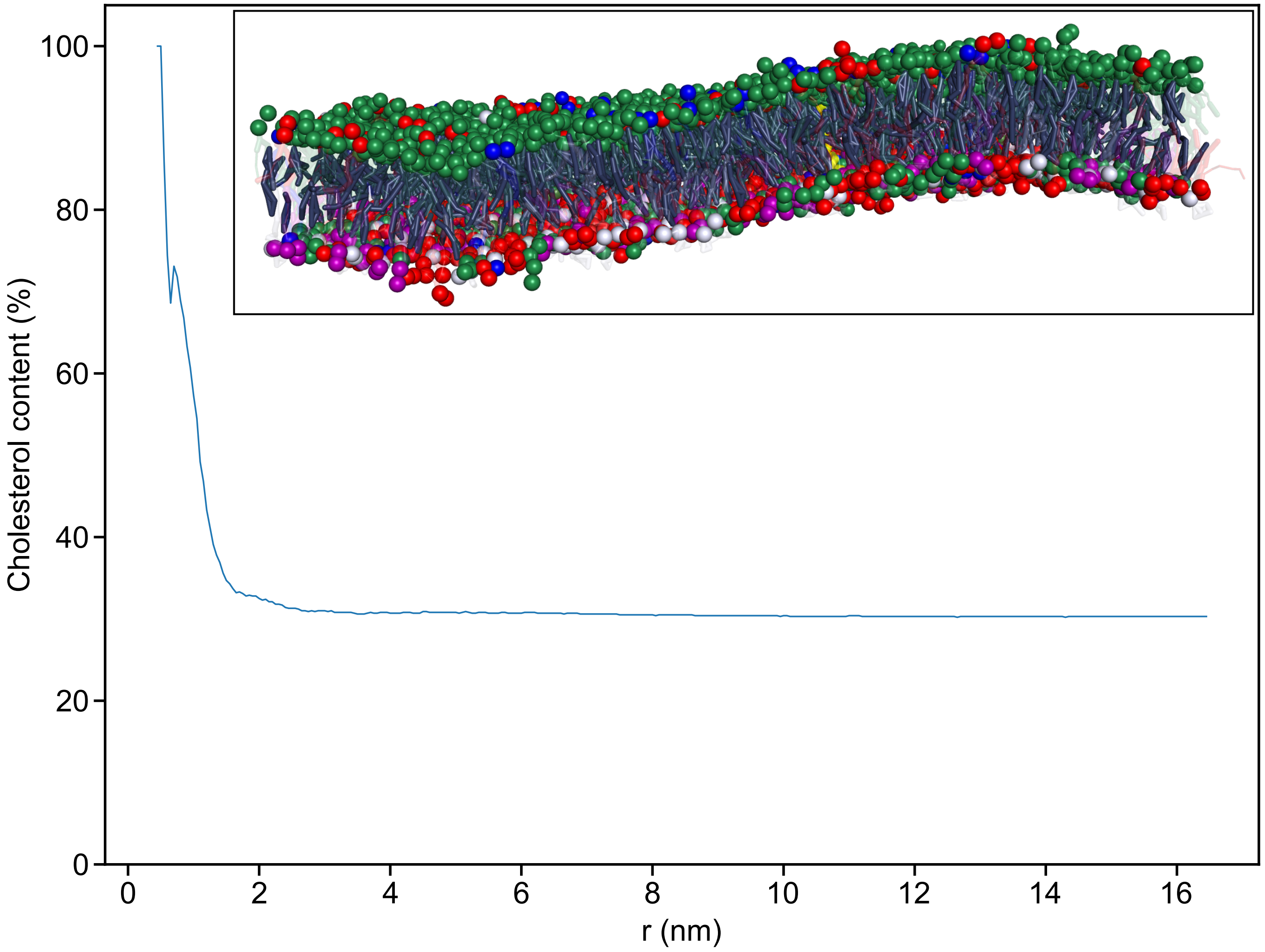


**Fig. S8.** A cholesterol attractor ($D_3K_3L_8K_3D_3$) produces high local cholesterol content (57.3% within 1 nm) in a native epithelial membrane model ((47), "Membrane 1").

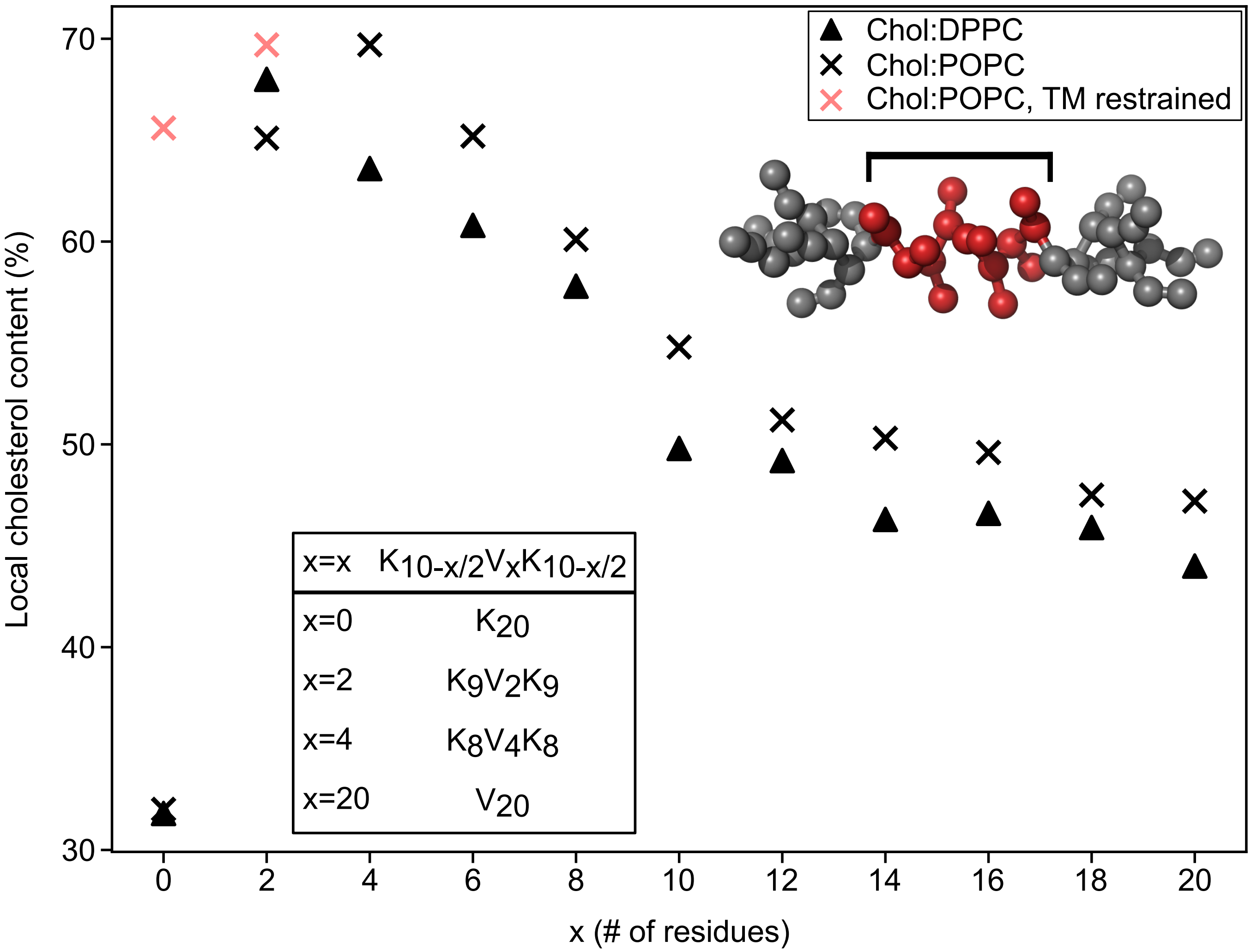


**Fig. S9.** Short, hydrophobic blocks produce a high local (1.0 nm radius) cholesterol composition of the membrane. Sequences adhere to the following motif: $K_{(10-x/2)}$–$V_x$–$K_{(10-x/2)}$ (x=0, 2, 4 etc.). The cholesterol attracting effect is present in both liquid-ordered (30% cholesterol, 70% DPPC) and liquid-disordered (30% cholesterol, 70% POPC) membrane phases. Restraining transmembrane position of very-short hydrophobic block sequences ($x \leq 2$) reveals high sensing functionality, persisting even in absence of the hydrophobic block ($K_{20}$).

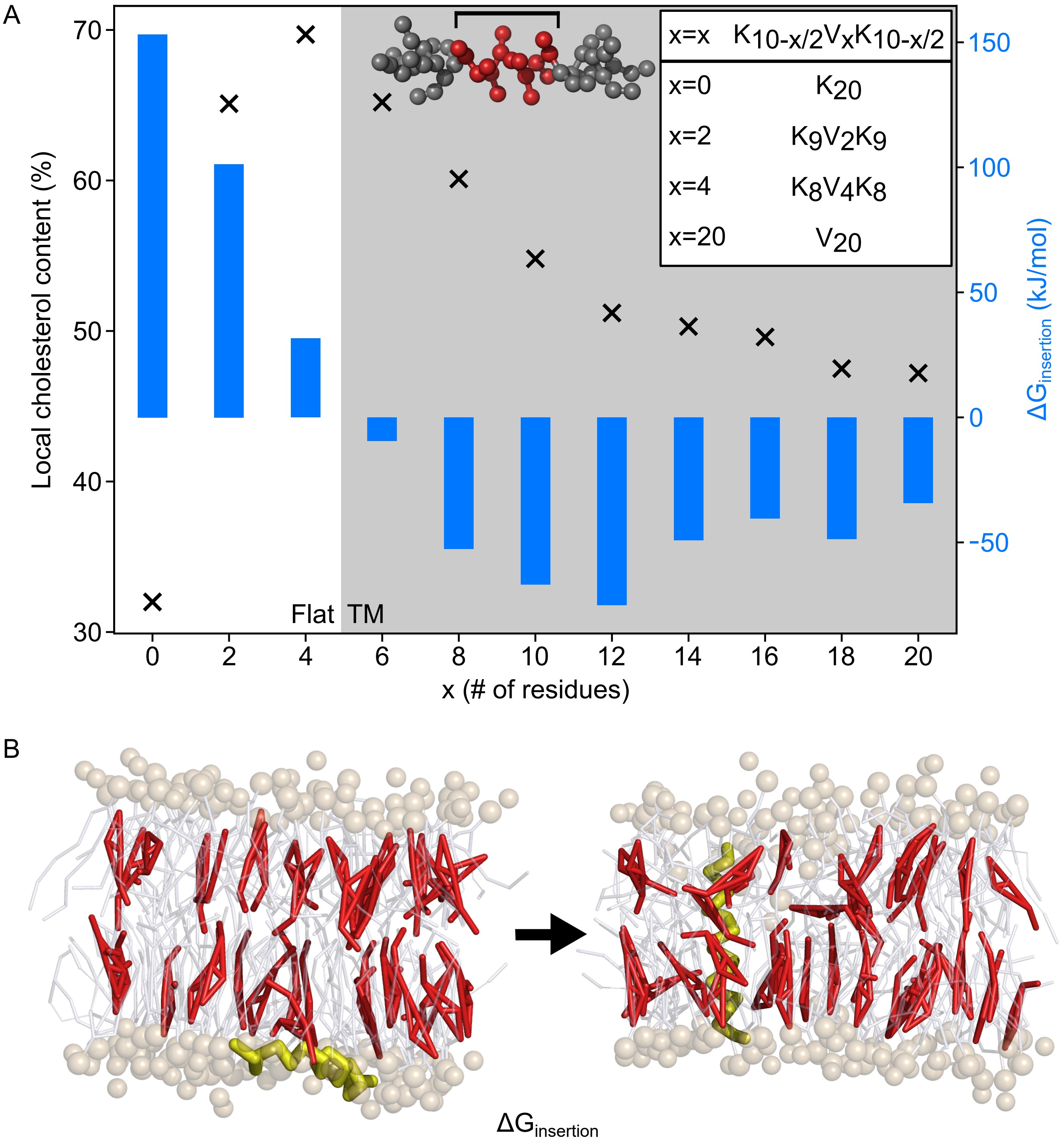


**Fig. S10.** The cholesterol sensing pattern follows from a trade-off between short hydrophobic block and stability of the TMD. (A): Sequences adhere to the following motif: $K_{(10-x/2)}$–$V_x$–$K_{(10-x/2)}$ (x=0, 2, 4 etc.). Cholesterol attraction prefers short hydrophobic blocks, while the TMD becomes unstable at very short blocks ($x \leq 4$). Optimal cholesterol attractors are therefore likely found in the TM-favoring region near metastability ($\Delta G = 0$), as is observed in the GA-determined sequence logo (6-8 hydrophobic residues). (B): TM stability of the peptide is judged based on the free energy of insertion, computed between state 1 (left: peptide (yellow) flat on membrane) and state 2 (right: peptide positioned within the membrane).

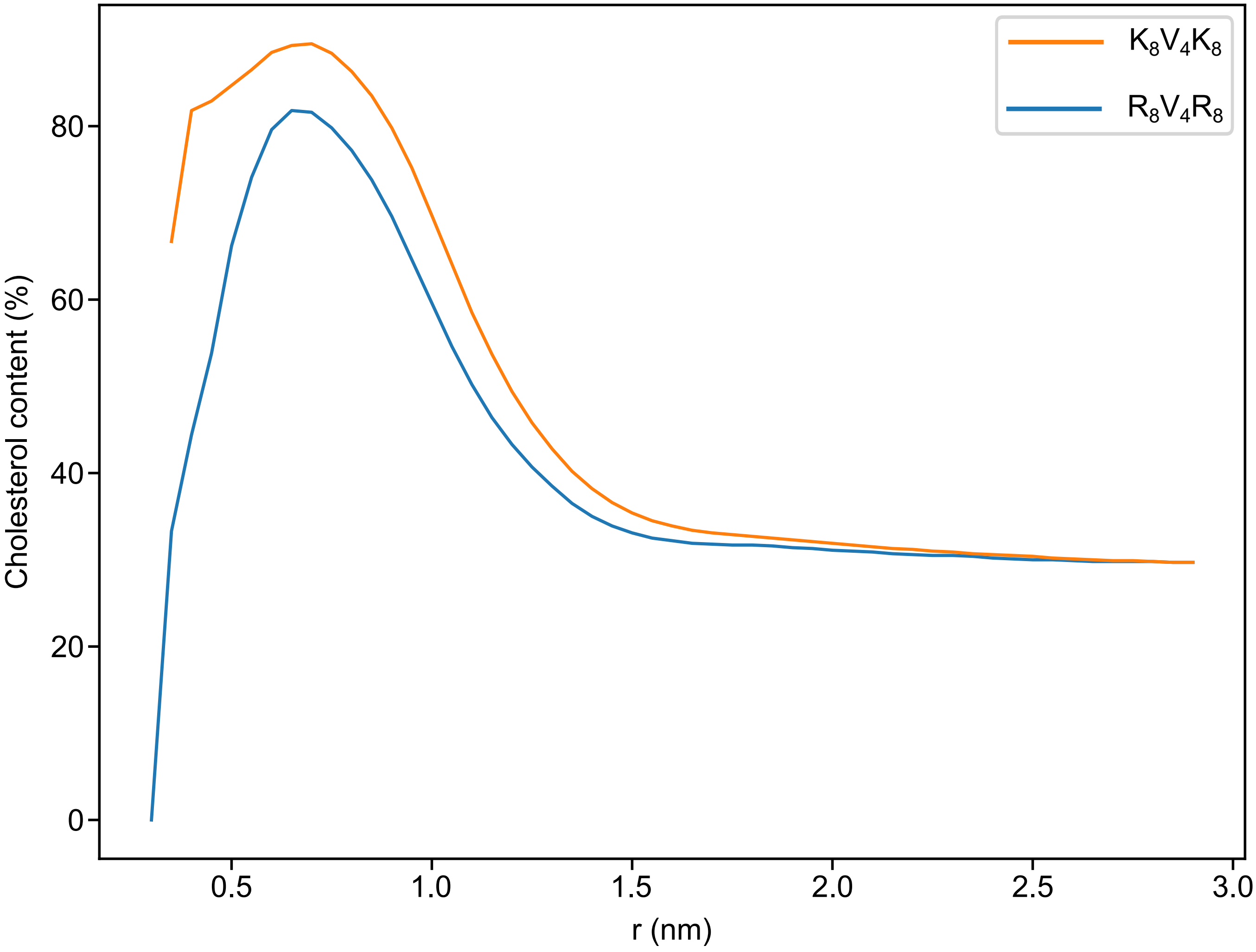


**Fig. S11.** Substitution of lysine (K) residues with arginine (R) residues in a dummy peptide sequence ($X_8V_4X_8$ (X=K,R)) leads to a small reduction in local membrane cholesterol content (lysine: 69.7% within 1 nm; arginine: 59.6% within 1 nm).

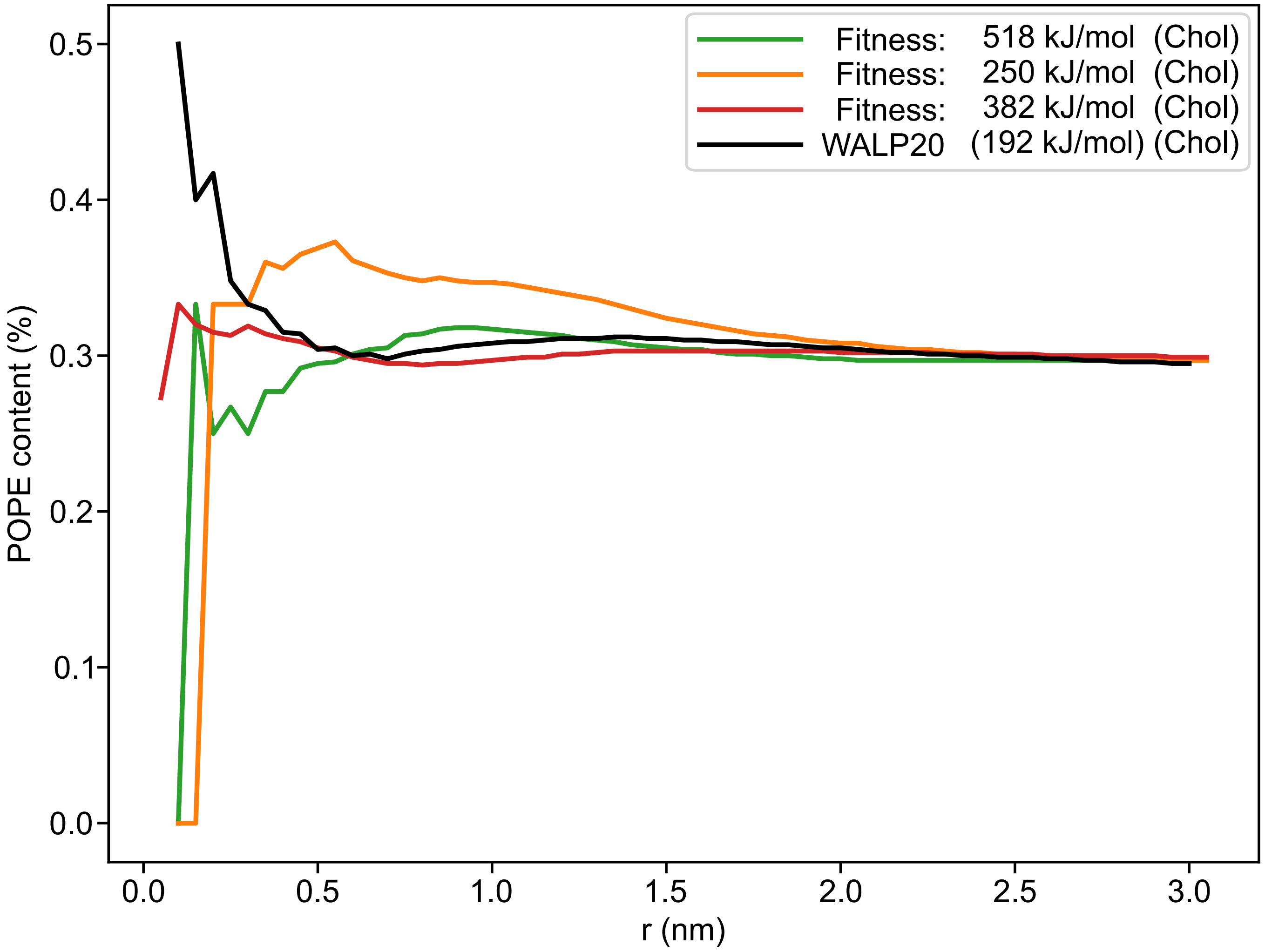


**Fig. S12.** The functionality of GA generated cholesterol attractors does not simultaneously correlate/translate to an increased affinity for POPE lipids in POPE/POPC (30%:70%) membranes. Ironically, the control peptide WALP20 turns out to be the best PE attractor. This illustrates that the attraction of cholesterol is mediated by a very different driving force than POPE attraction.

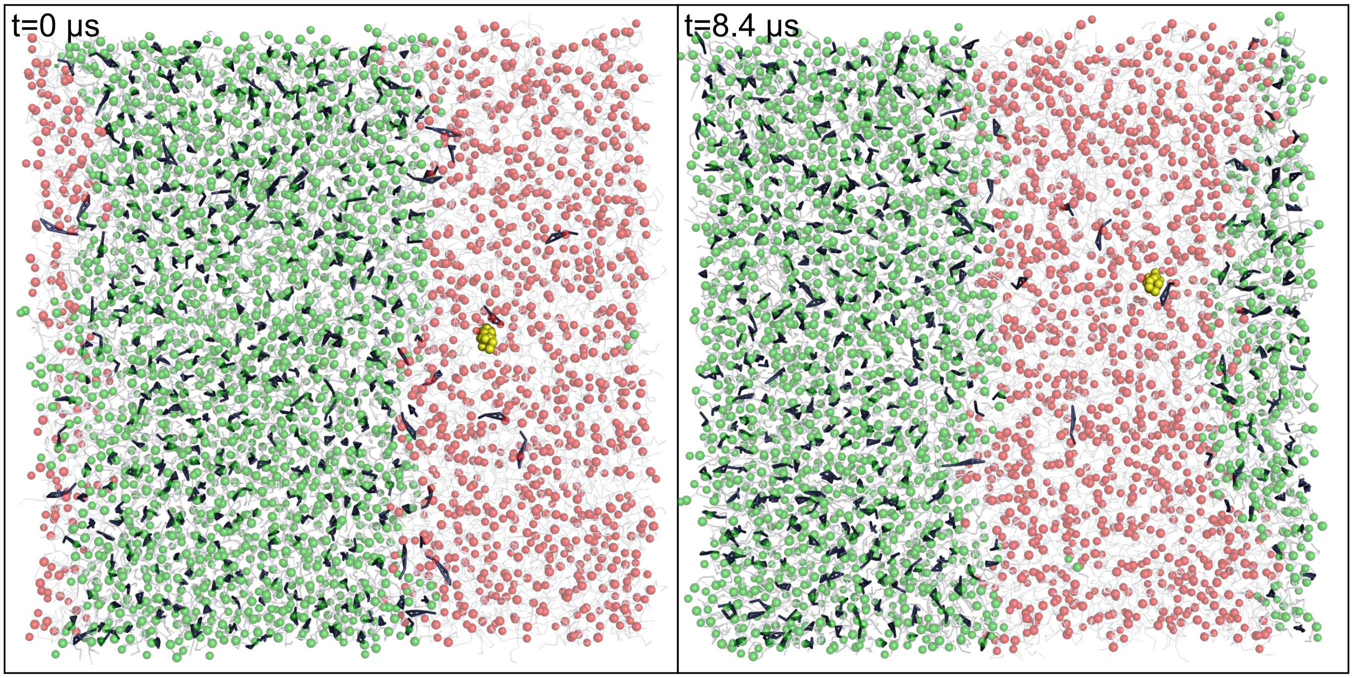


**Fig. S13.** A cholesterol attractor ($D_3K_3L_8K_3D_3$, in yellow) does not interact with the liquid-ordered/liquid-disordered (green/red) interface in a DPPC:DAPC:CHOL (50%:30%:20%) system.

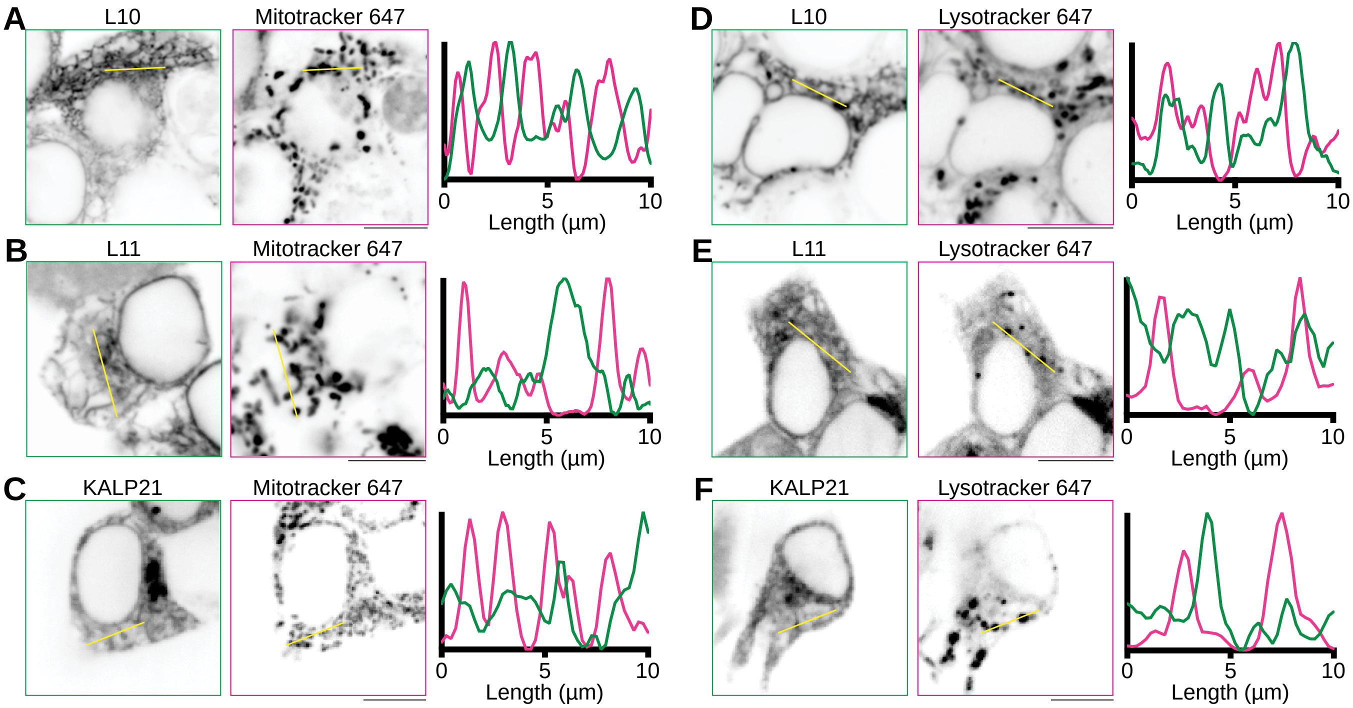


**Fig. S14.** L10 (A,D), L11 (B,E), and KALP21 (C,F) are distinct from the mitochondria (A,B,C) and lysosomes (D,E,F) in transfected HEK cells. For each panel, a line profile was drawn (yellow), and the normalized fluorescence intensity profiles of peptide and Mitotracker/Lysotracker are compared in the respective graphs. Scale bars and line profiles in all panels correspond to 10 μm.

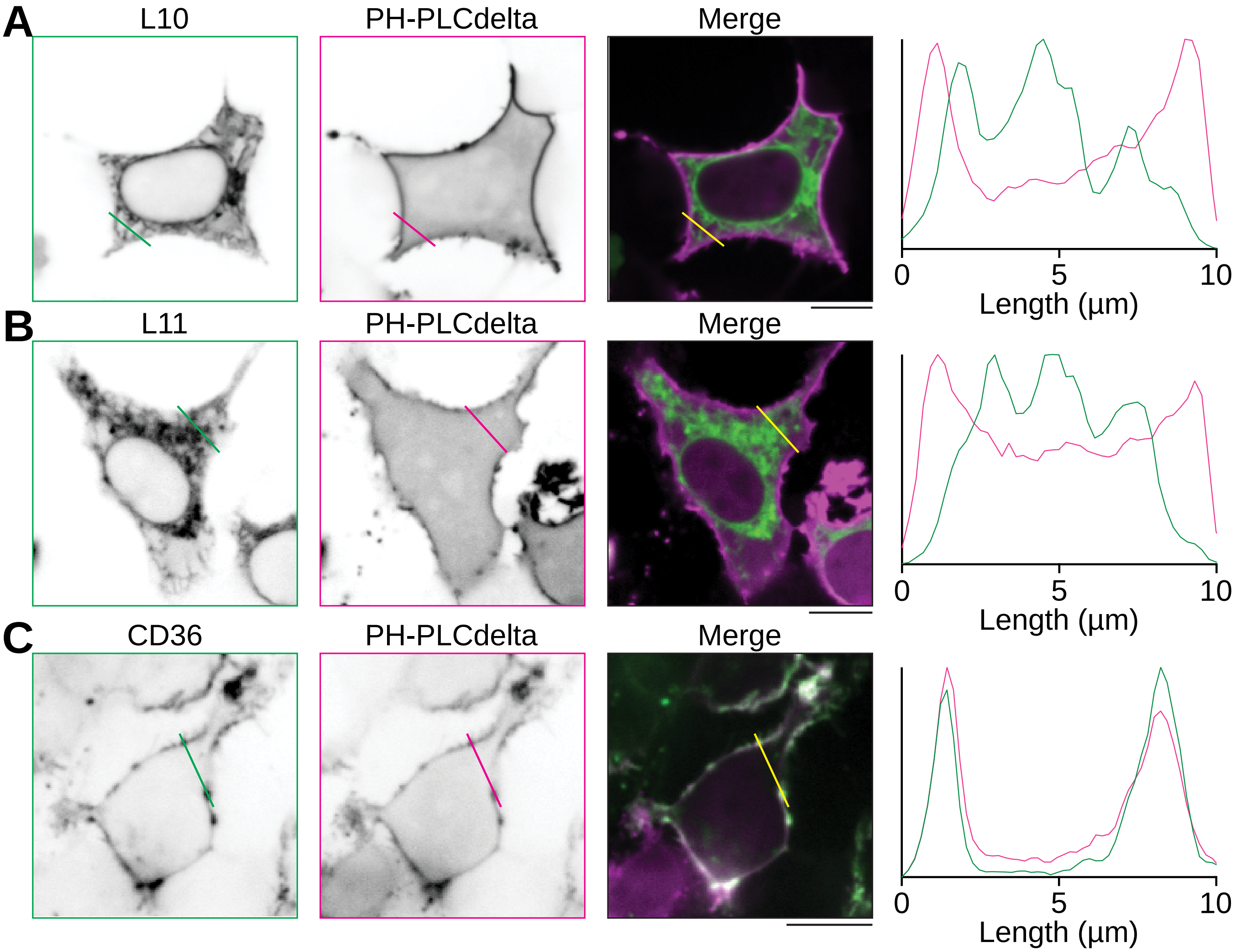


**Fig. S15.** L10 (A) and L11 (B) are distinct from the plasma membrane, marked using PH-PLCdelta (soluble membrane protein). (C): CD36 localizes to the plasma membrane. Scale bars and line profiles in all panels correspond to 10 µm.

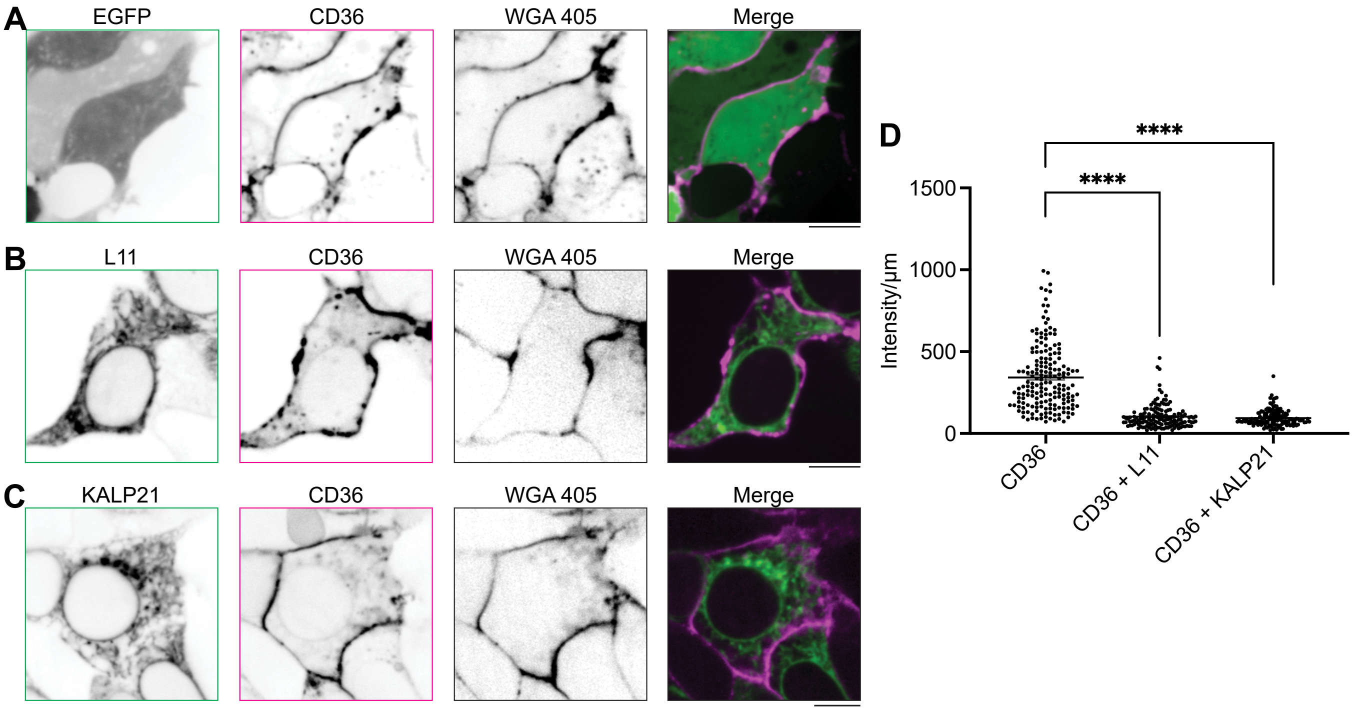


**Fig. S16.** L11 and KALP21 decrease the trafficking of fat transporter and scavenger receptor CD36 to the plasma membrane. (A) Control expression of soluble EGFP and mCherry-CD36 show the major fraction of CD36 in the plasma membrane in comparison with co-expression of mCherry-CD36 with either (B) L11 or (C) KALP21 peptides. Scale bars, 10 µm. (D) Quantification of the average fluorescence intensity of CD36 along the plasma membrane using WGA-405 as a plasma membrane marker. (Data from three independent experiments, ****$p<0.0001$, unpaired student t-test).
